# Beyond ERCs: exploring catastrophic forms of rDNA instability in aging yeast

**DOI:** 10.64898/2026.04.21.719800

**Authors:** Joseph O. Armstrong, Elizabeth X. Kwan, Gina M. Alvino, M. K. Raghuraman, Maitreya J. Dunham, Bonita J. Brewer

## Abstract

Single-celled organisms grown in identical conditions have variable life spans. Identifying the factors that drive the inherent variability in life span is crucial for our understanding of aging at a fundamental level. Here, we revisit the role of chromosome XII instability as a source of life span variability in aging populations of the budding yeast, *Saccharomyces cerevisiae*. We followed populations of mother cells as they aged and quantified changes in karyotype, DNA content, and aberrant structures within the rDNA. We quantified the kinetics of rDNA amplification both as extrachromosomal rDNA circles (ERCs) and as structural forms that are not resolved on CHEF gels. We propose mechanistic models of rDNA instability that describe how both ERCS and other aberrant structures are generated and the unique affects they have on life span. One of the models, which we call Catastrophic IntraChromosomal Repair (CICR), follows the fate of a broken replication fork in the rDNA locus when the template used for repair has already been replicated. At the completion of replication, when all other replication forks have successfully terminated, single, unopposed replication forks remain in the rDNA generating branched DNA molecules that have profound consequences during mitosis and/or subsequent cycles. In addition to ERCs, CICR and related events are additional forms of rDNA instability that contribute to the life span variability in isogenic populations of aging yeast.

**AUTHOR SUMMARY:** Aging is an uphill battle against entropy. The wear and tear on biological processes in cells require surveillance mechanisms and repair processes to stave off the inevitable. A cell’s genome is constantly under external and internal insults that can ultimately lead to death. Known events that impact a cell’s DNA are damage from free radicals (produced by the very oxygen needed for survival), shortening of the caps on chromosomes (called telomeres) every time a cell divides, and mutations that accumulate over time that affect essential cell processes. Yeast is an excellent tool to examine specific causes of aging because of its asymmetric mode of cell division: a mother cell can divide a limited, but variable number of times before senescing. We have isolated cohorts of mother cells, examined their genomes over the course of their life span and propose a new intrinsic insult on the genome. We find, in addition to the accumulation of extrachromosomal copies of the genes that encode the RNA portion of ribosomes (rDNA), that cells accumulate toxic runaway replication forks that can results in a branched and/or fragmented chromosome that leads to cell death.

## INTRODUCTION

Replicative life span (RLS) in the budding yeast *Saccharomyces cerevisiae* is defined as the number of daughter cells a mother cell can produce before senescing (Mortimer and Johnson, 1959). Life span is under genetic control as specific mutations significantly increase or decrease the median life span of a population; however, within an isogenic population, the exact number of times an individual cell divides is remarkably variable. What drives the inherent variability in life span? An attractive hypothesis explaining the variability of life span among isogenic individuals was first proposed by Sinclair and Guarente (1997) and centers on the observation that aging cells accumulate additional rDNA copies in the form of extrachromosomal rDNA circles (ERCs). Once formed, ERCs may replicate but are not free to segregate into daughter cells leading to a build-up of ERCs in aging mothers (Sinclair and Guarente, 1997; Shcheprova et al., 2008; Denoth Lippuner et al., 2014). The nature of ERC formation and subsequent accumulation creates a situation where the timing of an ERC’s formation is an important moment in the life of an individual yeast cell. A cell that forms an ERC early in life will accumulate more copies than one that forms one late in life, thus rDNA instability generates life span heterogeneity in an otherwise isogenic population.

How does rDNA accumulation lead to cell death? While the exact mechanism appears undecided, proposed mechanisms include titration of replication or transcription factors (Sinclair and Guarente, 1997; Falcón and Aris, 2005), sequestration of Sir2 and further rDNA destabilization (Yamamuro et al., 2025), excessive rDNA transcription that disrupts nuclear homeostasis (Morlot et al., 2019), remodeling of nuclear pore complexes (Meinema et al., 2022) and deregulation of the G1/S transition (Neurohr et al., 2018). Mutations in genes involved in the replication and maintenance of the rDNA locus (such as *fob1Δ* and *sir2Δ*; Defossez et al., 1999; Kaeberlein et al., 1999, respectively) can alter rDNA stability, ERC accumulation and ultimately the median life span of a population.

While ERCs are generally regarded to be the fatal product of rDNA instability, Takehiko Kobayashi (2008) proposed a broader rDNA theory of aging in which instability within the rDNA array itself drives senescence, not necessarily ERC accumulation. His proposal was supported by the finding that strains with reduced replication initiation activity have low levels of ERCs but nevertheless showed a reduced lifespan (Ganley et al., 2009). Additionally, the amplification of a region of chromosome XII telomere-proximal to the rDNA (Chr XIIr) was found in aging haploids (Hu et al., 2013) and diploids (Zylstra et al., 2023). Life span and rDNA instability are interconnected, but the relationship between the rate of ERC accumulation and lifespan is not a simple one.

There are multiple aspects of the structure, replication and transcription of the yeast rDNA locus that contribute to its inherent instability. First, the tandemly repeated nature of the rDNA locus (Fig 1A) makes it prone to copy number changes at a much higher rate than the rest of the genome (S1A,B Fig; Szostak & Wu, 1980; Petes, 1980; Kobayashi, 2011; Kwan et al., 2016). Second, it is the most highly transcribed region of the yeast genome (Warner, 1999) even though only about half of the repeats are active at any given time (Dammann et al., 1993) and origins downstream of actively transcribed repeats are preferentially activated for replication (Fig 1B; Pasero, et al., 2002). Third, in the absence of Sir2, PolII transcription from the intergenic spacer (*NTS1*; Smith and Boeke, 1997; Fritze et al., 1997; Li et al., 2006) displaces cohesin, increasing the frequency of unequal sister chromatid recombination in the rDNA (Gottlieb and Esposito, 1989; Kobayashi et al., 2004; Kobayashi and Ganley, 2005). Fourth, while the rest of the genome is replicated by bidirectionally oriented forks, the fates of the two forks initiated in the rDNA are not equivalent as most of the replication proceeds unidirectional by a fork moving in the direction of 35S transcription (Fig 1C; Brewer and Fangman, 1988; Linskens and Huberman, 1988). The oppositely oriented fork attempting to enter the 3’ end of the adjacent 35S transcription unit is arrested at a site called the replication fork barrier (RFB; Brewer and Fangman, 1988) by the binding of the Fob1 protein (Fig 1B,C; Kobayashi and Horiuchi, 1996) where it awaits the arrival of the adjacent, oppositely oriented replication fork.

**Figure 1:**
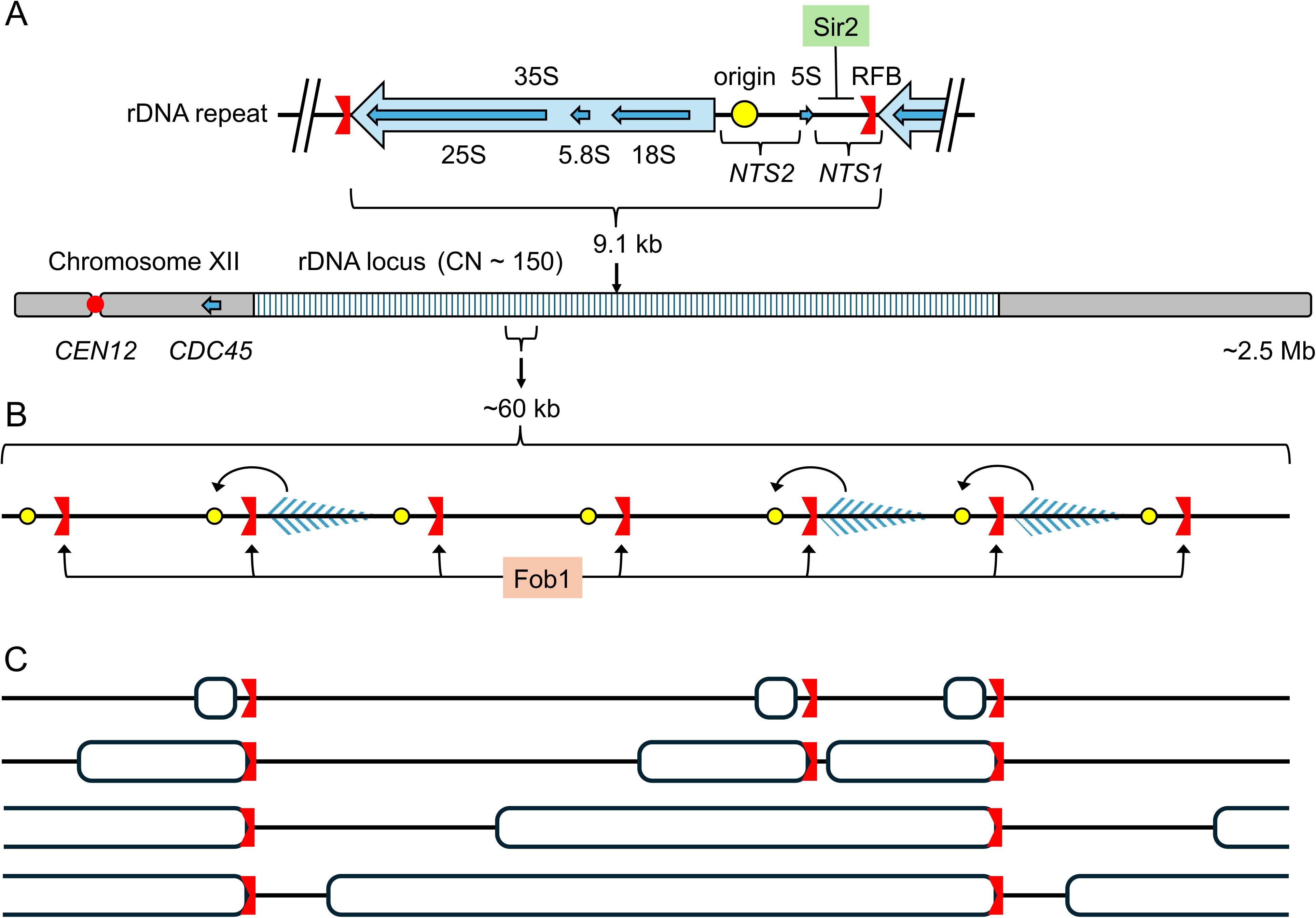
Structure, transcription, and replication of the rDNA locus on Chr XII. A) The rDNA locus is found on the 2.5 Mb Chr XII and is composed of ∼150 copies of the 9.1 kb rDNA repeating unit. Each unit is identical in sequence and encodes two transcripts (35S and 5S), a potential origin of replication, and a replication fork barrier (RFB). The <u>n</u>on<u>t</u>ranscribed spacers (*NTS1* and *2*) contain multiple regulatory sequences, including a PolII transcript repressed by the presence of Sir2. *NTS2* and the unique gene *CDC45* were used as hybridization probes in this work. B) 35S transcription is restricted to roughly half of the repeats that are devoid of nucleosomes (Dammann et al., 1993). The origins downstream of actively transcribed units are preferentially activated for replication (curved arrows; Muller et al., 2000; Pasero et al., 2002). Fob1 binds to the replication fork barrier (RFB) to prevent head-on collisions between the rightward moving replisome and the leftward moving RNA polymerases in the 35S transcription unit. C) Replication of this section of chromosome is illustrated as a series of drawings at progressively later times in S phase. The RFB/Fob1 complex enforces the mostly unidirectional replication of this large region of Chr XII (Kobayashi and Horiuchi, 1996) with forks moving leftward toward the centromere.

An rDNA instability event can be initiated when the arrested fork at the RFB breaks, resulting in a single-ended, double-stranded break at the RFB (Kobayashi et al., 1998). In general, a broken replication fork can be repaired in two ways (Fig 2A): by break induced replication (BIR; Fig 2Ai-iv; Saini, et al., 2013; Kramara et al., 2018; Qi et al., 2026) which establishes a conservative form of replication with a migrating D-loop or by replication fork restoration (RFR; Fig 2Av-vi; Amiama-Roig et al., 2025) which restores the semiconservative replication fork. While progression of either of these forks would be halted at the adjacent RFB, either fork allows normal termination to occur when met by a fork from the adjacent replicon.

**Figure 2:**
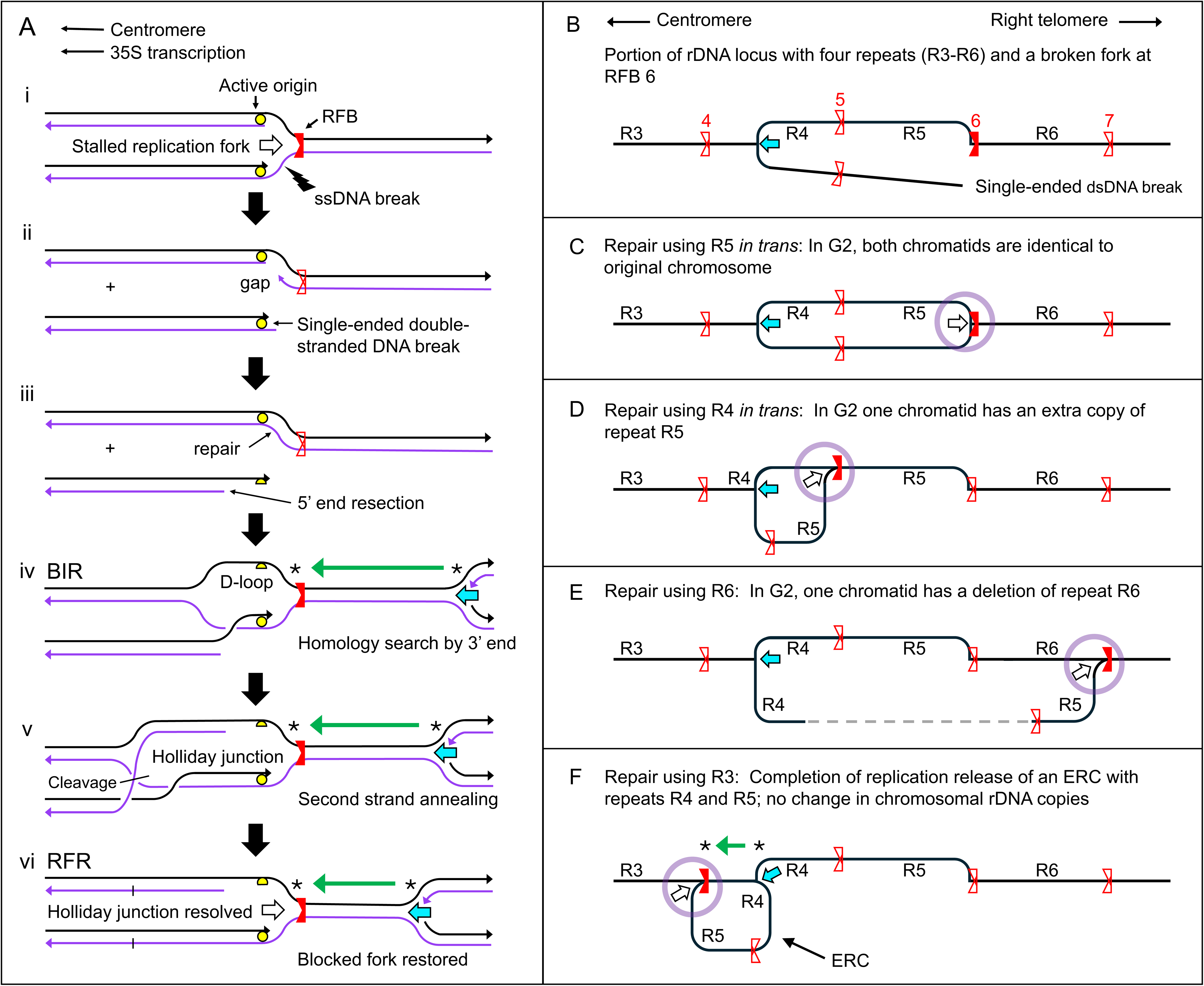
Pathways for the repair of single-ended, double-strand breaks at the rDNA replication fork barrier. A) The break of an arrested replication fork produces a single ended, double-stranded break (i and ii). After 5’ end resection (iii) the 3’ single stranded end begins a homology search (iv). This invasion can lead to break induced replication (BIR) in which the invading strand is extended by conservative replication as the D-loop migrates rightward. However, in this case the fork is immediately arrested at the RFB. Instead, if the second strand invades the other strand in the D loop a single Holliday junction results (v). After resolution of the Holliday junction, the replication fork is restored but again is immediately arrested at the RFB (vi). Either BIR or RFR forks can serve as the termination site for advancing forks from the adjacent replicon (green arrow). B) Four repeats in the rDNA locus are shown with a single ended dsDNA break at RFB6. The dsDNA is now shown as a single line. Repair in the subsequent images occurs as described in A. C) If the single-ended dsDNA break uses the original repeat (R5) as a repair template, the original fork is restored and the two chromatids will have the same number of rDNA repeats. D) If repeat 4 is used for repair one chromatid will have an extra copy of R5. E) If repeat 6 is used for repair one chromatid will be missing R6. F) Repair using repeat 3 will generate an ERC when the two forks meet at RFB4 (green arrow). The two sister chromatids will be unchanged.

The repeat used for repair of a broken fork is usually illustrated as being the sister strand at the site of the break (Fig 2A-C); however, repair could also use a nearby un-replicated repeat as a repair template (Fig 2D-F; Ganley et al., 2009). These alternate choices for repair template would result in gain (Fig 2D) or loss (Fig 2E) of a repeat in the rDNA locus of one of the sister chromatids or in the formation of an ERC (Fig 2F) with no change in repeat number in either sister chromatid. The phenotypes of a *fob1Δ* mutant—longer RLS, no fork arrest at the RFB, a more stable chromosomal rDNA copy number (S1 Fig) and fewer ERCs (Kobayashi et al., 1996; Kobayashi et al., 1998; Defossez et al., 1999)—support the hypothesis that replication fork breakage and repair contribute to aging in yeast.

To characterize the kinetics and dynamics of rDNA instability across replicative lifespan we aged populations of wild type, *fob1Δ* and *sir2Δ* cells and evaluated their karyotype with Contour-clamped Homogeneous Electric Field (CHEF) gel electrophoresis, Southern blotting and flow cytometry. We found that ERCs accumulated in an age and genotype dependent manner, but the dominant form of rDNA instability we observed failed to migrate during CHEF gel electrophoresis. We propose that these species represent a distinct form of rDNA instability, whose product contains unopposed replication forks that hinder migration in gel electrophoresis. We generated two models to describe how these unopposed replication forks can occur.

Both models highlight the important role that the choice of template plays in determining the outcome of break repair. When these events occur in the chromosomal rDNA array (in a process which we call Catastrophic IntraChromatid Repair (CICR)), the product is a branched form of Chr XII. When this process occurs in replicating ERCs, the product is a multiply-branched, double-lariat structure (DL-ERC). We propose that these events, along with ERC accumulation, contribute to lifespan variation within isogenic populations.

## RESULTS

### Generating populations of aging mother cells

We used miniature aging chemostat devices (MADs, Fig 3A; Hendrickson et al., 2018) to cultivate large populations of aging mother cells of wild type, *fob1Δ* and *sir2Δ* strains. We inoculated two chemostats on day 0 with cells from an overnight culture. Over the next six days, mother cells were continuously supplied with excess medium as the daughter cells were removed in the effluent. We periodically assessed viability of mother cells using a membrane permeable stain and flow cytometry (n > 10,000, Fig 3B) and we quantified the replicative age of the populations over time by staining and manually counting bud scars (n = 40, Fig 3C). We observed that *fob1Δ* mother cells accumulated the highest average number of bud scars (mean: 19.4) and maintained viability the longest of the three populations, while *sir2Δ* mother cells accumulated the lowest average bud scars (mean 10.4) and exhibited more rapid loss in viability. These results align with previously reported effects of *FOB1* and *SIR2* deletions on life span (Defossez et al, 1999; Kaeberlein et al., 1999). The maximum replicative age observed for each genotype occurred on different days (*fob1Δ*, day 6; WT, day 4; *sir2*Δ, day 3; Fig 3C). The mean replicative age of the wild type and *sir2*Δ populations slightly decreased after reaching a maximum, likely due to differences in cell cycle progression among individual cells.

**Figure 3:**
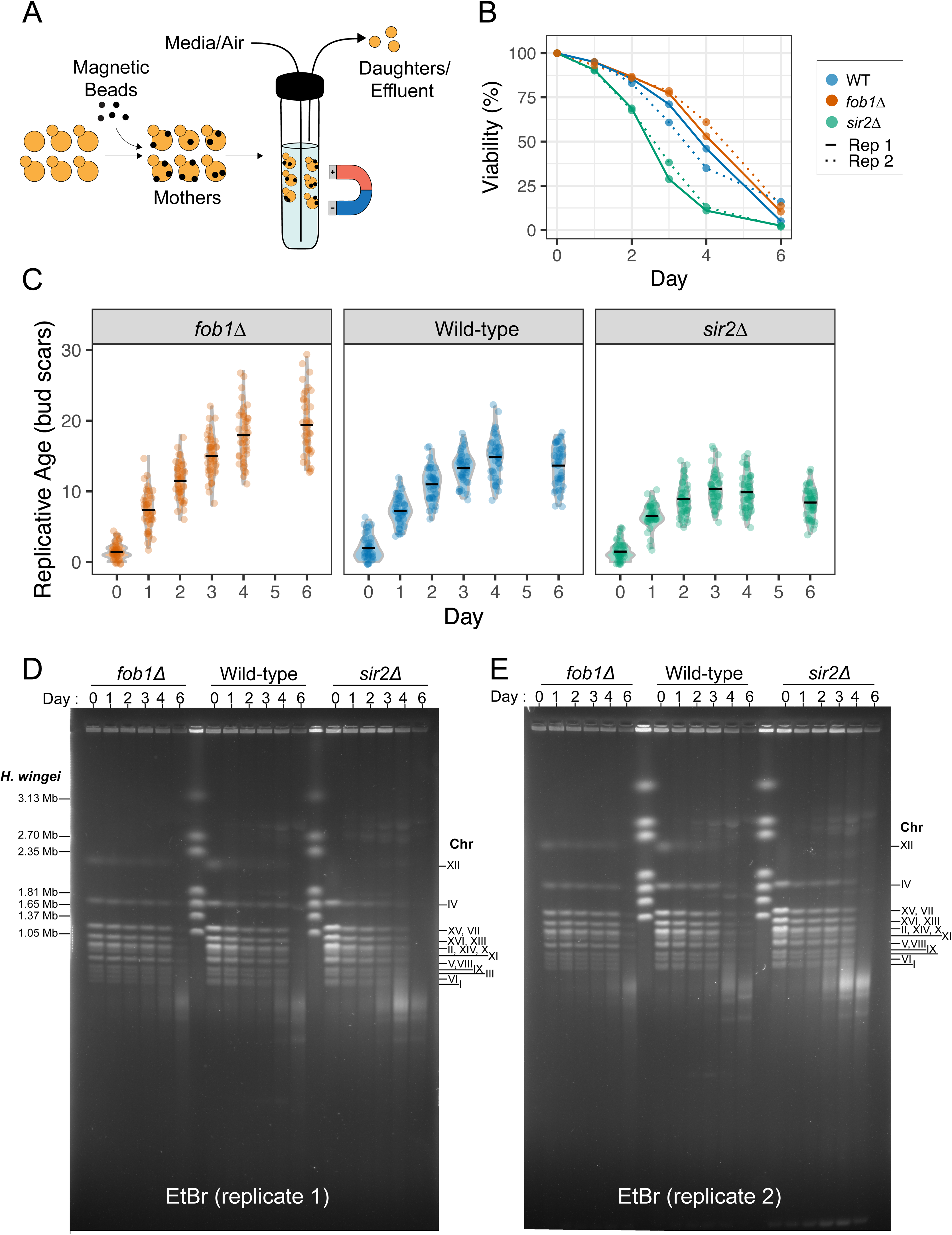
Karyotyping the genome from aging chemostats using CHEF gel electrophoresis. A) Miniature aging chemostats were used to isolate large populations of aging mother cells. Magnetic beads were covalently bound to mother cells which were retained on the chemostat wall by an external magnet while being continuously perfused with fresh media to maintain a constant environment and to remove daughter cells. B) Viability of mother cells was assessed with propidium iodide and flow cytometry (n > 10,000). Dashed lines represent the second replicate population. C) Replicative age was determined by manually counting bud scars of one mother population for each genotype over time (n = 40 for each day). The horizontal bar indicates the average number of bud scars for the populations. D and E) Ethidium bromide-stained CHEF gels made from mother cells sampled across six days of growth from the two replicate populations of each of the three genotypes. Individual bands contain one or more comigrating chromosomes. As cells age, additional DNA species become apparent.

### Karyotyping aging genomes

We collected samples from the chemostats over the replicative life span of each population and prepared agarose plugs for karyotyping by CHEF gel electrophoresis. The ethidium bromide-stained gels of two replicate sets of aging mother populations are shown in Fig 3D,E. Most of the bands in the CHEF gels contain one or more comigrating chromosomes, with Chr XII being the largest and least mobile. By day 1 Chr XII disappeared from its normal location in the gel and additional DNA species both above and below the position of Chr XII appeared.

To identify individual chromosomes, we hybridized Southern blots of the two CHEF gels with unique probes from Chrs XII, IV, X and VIII (*CDC45*, *GAL3*, *ILM1*, and *CUP1,* respectively; replicate 1, Fig 4A-D and replicate 2, S2A-D Fig). There were three regions of the CHEF gels with hybridizing material—the well, full-length linear chromosomes (at unique positions for each chromosome) and a smear of chromosome fragments that was in the same size range for all four probes. On day 0, Chrs VIII, X and IV migrated in the gel at positions consistent with their linear sizes. Chr XII appears as a discrete band for wild-type and *fob1Δ* populations at a size that is consistent with ∼125 rDNA copies per chromosome (Fig 4A and S2A Fig). However, for the *sir2Δ* mutant, Chr XII appears as a broad distribution of sizes reflecting the variability in chromosomal rDNA repeat number. We observe a slight increase in chromosomal rDNA copy number among clones derived from day 6 (old) WT and *sir2Δ* mother populations compared to the starting population but not for *fob1Δ* populations (S1B,C Fig). We also observe a slight increase in the coefficient of variation when day 6 WT and *sir2Δ* clones were compared to clones derived from glycerol stocks (young, S1D Fig).

**Figure 4:**
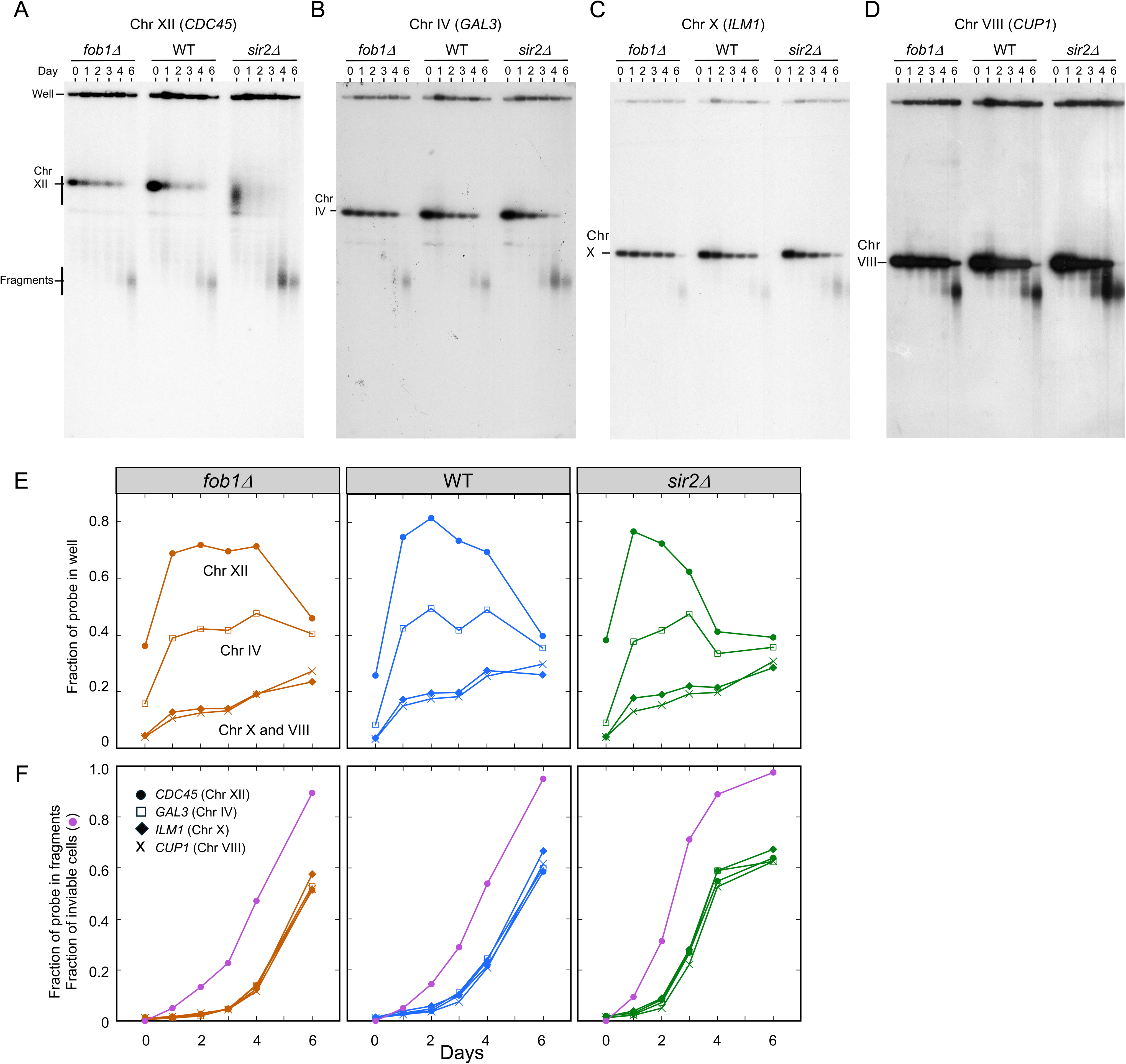
Characterizing the distribution of individual chromosomes in CHEF gels. A-D) The autoradiograms of the CHEF gel shown in Fig 3E (replicate 1) were produced by Southern hybridization using fragments from *CDC45*, *GAL3, ILM1* and *CUP1* to identify the positions in the gel of Chrs XII, Chr IV, Chr X and Chr VIII, respectively. E and F) The total amount of hybridization in each lane was used to calculate the fraction of each probe remaining in the well (E) and in the fragments (F). These values are estimates of the fraction of each chromosome that remains in the well or that has been fragmented over the six days of chemostat growth. The kinetics of inviability is also plotted in F. Similar data for replicate 2 can be found in S2 Fig.

By day 1 we found the amount of each chromosome retained in the well increased (Fig 4E and S2E Fig), likely due to the presence of DNA branches from on- going replication. The modest accumulation of DNA from the two smaller chromosomes (Chrs VIII and X) is consistent with the observation that these two chromosomes do not contain large regions of late replicating DNA (Foss et al., 2017; Alvino et al., 2007) and thus complete replication in a narrow S-phase window. This result sets an upper limit on how much of the well-retention could be due to inefficient lysis of the plugs. Completion of replication is expected to take longer for both Chr IV and XII due to the presence of regions of ∼300 kb that are late replicating (Alvino et al., 2007; Foss et al., 2017) and well-retention is increased for both chromosomes over that for Chr VIII and X (Fig 4E and S2E Fig). However, compared to Chr IV, very little intact chromosome XII migrated into the gel. This finding is consistent with the observation that the rDNA in Chr XII in aging cells is branched, causing Chr XII to remain in the well (Pal et al., 2018).

The smear of DNA in the size range of 50 to 200 kb hybridizes to all four unique probes and in similar proportions that track with the genotype of the strain (Fig 4F and S2F Fig). The accumulation of fragmented DNA follows the appearance of inviable cells and suggests that fragmentation of chromosomes is not the cause of cell death but rather the consequence of cell death.

The new bands above the position of Chr XII did not hybridize with the *CDC45* probe (Fig 4A; S2A Fig) and indicate that this new species of DNA is not a larger form of Chr XII. Using a probe from the non-transcribed portion of the rDNA repeat (*NTS2,* Fig 1A; Fig 5A; S3A Fig) we found that the DNA bands migrating both slower and faster than Chr XII contained rDNA with an overall increase in *NTS2* hybridization that is age and genotype specific. Could these slowly migrating species be ERCs?

**Figure 5:**
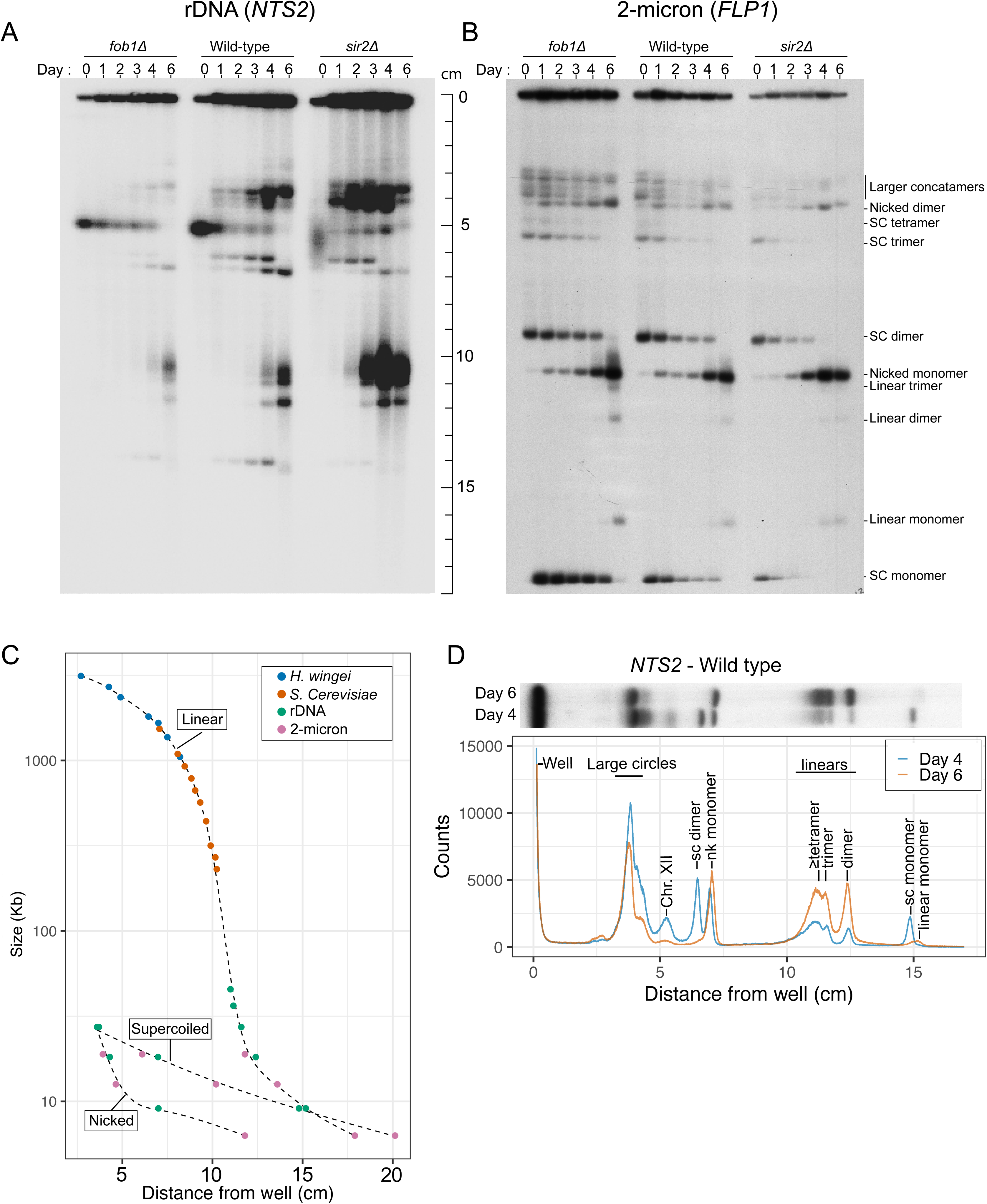
Identifying and characterizing ERC isomers. A and B) The autoradiograms of the CHEF gel shown in Fig 3E (replicate 1) were produced by Southern hybridization using probe fragments *NTS2* and *FLP1* to identify the positions in the gel of potential ERCs and the 2-micron plasmid, respectively. The linear forms of 2-micron were identified by their appearance on day six of each culture. Supercoiled (SC) forms, which migrate faster than linear forms in convention gels, were expected to disappear as they become converted to nicked circular (NC) forms as cells die over the aging time course. The nicked circular forms were identified by their reciprocal increase in intensity over time. C) The distances migrated (in cm) for the *S. cerevisiae* and *H. wingei* chromosomes and the linearized 2-micron multimers were used to establish a standard curve for linear DNA molecules. The distances migrated for nicked and supercoiled circular forms were plotted against their predicted mass to generate standard curves for supercoils and nicked circles. The intermingling of nicked or supercoiled forms of ERCs and 2-micron circles confirms the identity of topological and concatemeric forms of ERCs. D) Tracings of *NTS2* hybridization on days 4 and 6 of the wild type mothers reveals the relative amounts of the various forms of extrachromosomal rDNA. Between days 4 and 6 the supercoils have been converted to nicked and linear species. The *FLP1* and *NTS2* autoradiograms for replicate 2 of the aging experiment are shown in S3 Fig.

### Characterizing rDNA structural forms

To deconvolute the numerous bands present on the *NTS2* blots, we compared the mobilities of the presumptive ERCs to that of the endogenous 2-micron DNA plasmid (6.3 kb; Fig 5B, S3B Fig) that exists as monomer circles (supercoiled and nicked) and as concatemers and catenanes with variable numbers of repeats. While migration of linear molecules is largely affected by their mass, circular and branched species are particularly sensitive to electrophoresis conditions (voltage, pulse time/regime, and agarose concentrations; Simske and Scherer 1989, Khan and Kuzminov, 2013, Beverly, 1988). Supercoiled monomer circles are compact, while their nicked counter parts have an open conformation that can be temporarily and repeatedly impaled on agarose spikes, slowing down their migration. Despite the small size of the 2-micron plasmids, we found that multimeric forms of 2-micron migrated at rates similar to chromosomes more than 100 times their mass (Fig 5B; S3B Fig). Over the course of the 6 days, as DNA fragmentation occurs in increasing subsets of cells, we see decreases in some bands (supercoils) and a reciprocal increase in other sets of bands (nicked). On day 6 (Fig 5B; S3B Fig) we see the appearance of bands that correspond to linear forms of the different 2-micron concatemers. We see similar shifts in subsets of bands in the *NTS2* blots (Fig 5A; S3A Fig) allowing us to identify different forms of ERCs and to estimate the number of rDNA repeats each contains (Fig 5C,D). From this analysis we conclude that, in aging mother cells, the majority of the *NTS2* signal migrating in the gel corresponds to ERCs containing between one and five rDNA repeats.

### Quantifying age-related increases in rDNA species

Visual comparison of the *NTS2* and *CDC45* blots (Fig 4A and 5A; S2A and S3A Fig) indicates that, in addition to increases in ERCs with age, the rDNA signal is also increasing in the well and fragmented DNA. To quantify increases in rDNA copy number (well, chromosome, ERCs or fragments) we normalized the *NTS2* signals to the single copy, total Chr XII probe (*CDC45*: well + chromosome + fragments; Fig 6, top).

**Figure 6:**
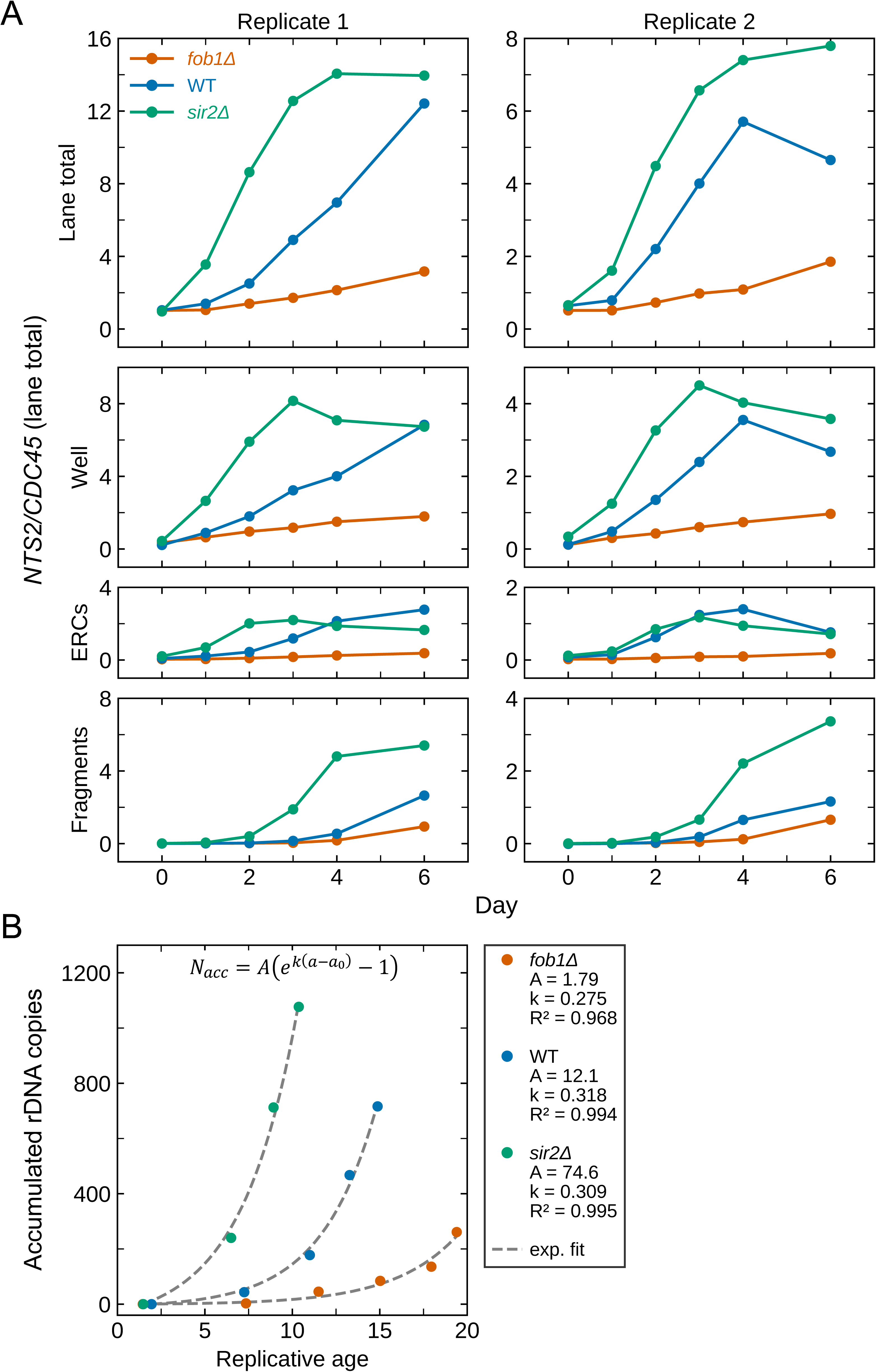
Estimating the increase in rDNA copy number during aging of the *fob1Δ*, WT and *sir2Δ* strains. A) The signal for each form of *NTS2* on days 0 through 6 was normalized to the total signal from *CDC45* in each gel lane. The total relative increases in rDNA repeats in replicates 1 and 2 are illustrated in the top panels. The *NTS2* signal remaining in the well, freely migrating ERCs and fragmented forms are shown in the bottom three panels. The differences in ratios on the Y-axes between replicates 1 and 2 are due to the difference in time that the blots were exposed to phosphor screens before scanning and quantification. B) Accumulated rDNA copy number was calculated using a fold change of the ratios in Fig 6A for each day relative to day 0, multiplied by the starting rDNA copy number (125 for WT and f*ob1*Δ, 90 for *sir2Δ*). Accumulated copies are total copies minus the starting copy number. Replicative age is the mean bud-scar count of mothers at each sampling time. Only samples where mean age increased from the previous day are shown (fob1Δ days 0-6, WT days 0-4, sir2Δ days 0-3). Dashed grey lines are the least squares fit to (*e^k^*^(*a*–*a*0)^ − 1) where *N_acc_* is the accumulated copies at a given age, a is the mean replicative age, a_0_ is the mean replicative age at day 0, *k* is the rate constant for rDNA accumulation and A reflect the frequency of an initiating event. The data shown are from replicate 1.

We found that the ratio of *NTS2* to *CDC45* increased between 11.9 - 14.4 fold compared to day 0 for the two replicate *sir2Δ* populations. The wild type populations (replicate 1 and 2) increased between 7.2 - 12 fold and the *fob1Δ* population increased 3.1 - 3.6 fold relative to day 0. The bulk (>50%) of the *NTS2* signal for all three strains did not migrate during electrophoresis and remained in the well (Fig 6A). The amount of signal present in the well and as mobile ERCs reached similar levels in WT and *sir2Δ* populations; however, the signal increase for wild type populations appeared roughly a day after the *sir2Δ* populations. The signal from fragmentated DNA also follows a similar pattern.

To quantify the kinetics of rDNA accumulation relative to mother cell age (bud scars) we calculated the total accumulated rDNA copies using the fold change in *NTS2/CDC45* ratio relative to day 0 and plotted total copies by mother cell age for replicate 1 (Fig. 6B). We fit the accumulated rDNA to the equation *N_acc_* = *A*(*e^k^*^(*a*–*a*0)^ − 1) where *N_acc_* is the accumulated copies at a given age, a is the mean replicative age, a_0_ is the mean replicative age at day 0, *k* is the rate constant for rDNA accumulation and A reflect the frequency of an initiating event. We found that values for *k* only differed ∼16% for the three populations (k = 0.275, 0.318 and 0.309 per division) indicating that it takes 2.52, 2.18 and 2.24 divisions (for *fob1Δ,* WT and *sir2Δ* populations, respectively) for the extra rDNA to double. The similarity of *k* among the three populations suggests that once an rDNA instability event occurs, rDNA copies increase at similar rates for each genotype. While values for *k* are relatively similar between genotypes, values for A varied substantially. We used A/A_wt_ ratio to calculate a relative probability of rDNA accumulation among strains and find that it varies from 0.15 for *fob1Δ* populations to 7.2 for *sir2Δ* populations, indicating that *fob1Δ* cells are roughly 7 times less likely to accumulate rDNA than WT populations and *sir2Δ* cells are roughly 7 times more likely than WT to accumulate rDNA.

The identification of migrating ERCs is unambiguous, however, the signal in the well is a composite of at least two forms of *NTS2*—replicating Chr XII and replicating ERCs. We estimated the contribution of chromosomal rDNA copies in each sample by relying on the *NTS2*/*CDC45* ratio in the well on day 0, when cells have no significant ERC population. Even though cell number decreased on subsequent days, we used the *CDC45* signal to calculate the amount of *NTS2* signal in the well that was due to full-length Chr XII (Fig 7A,C; rust, blue and green bars).

**Figure 7:**
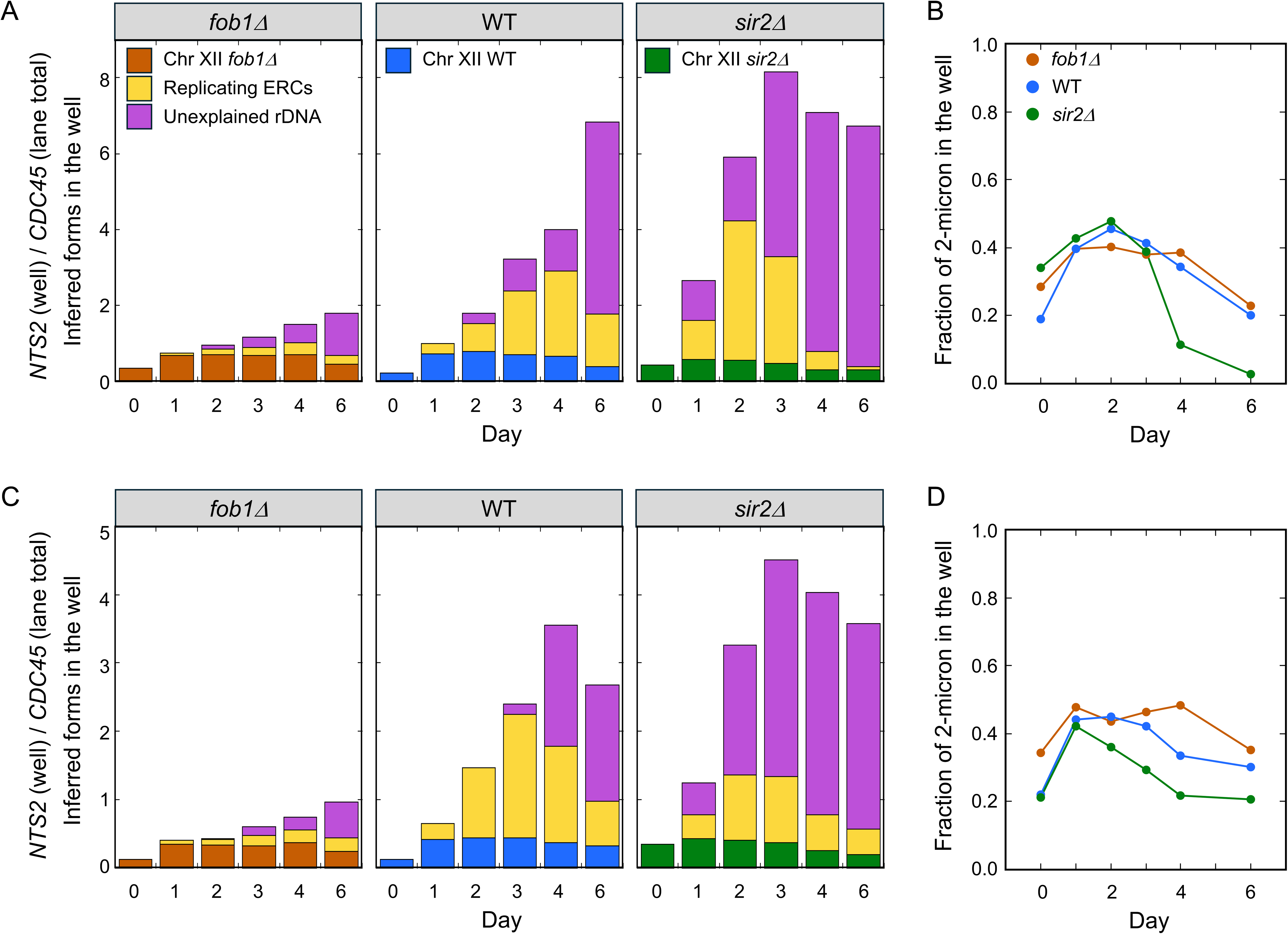
Analyzing non-mobile rDNA in CHEF gel. A and C) For each day, the total signal intensity of *NTS2* (rDNA) and *CDC45* (Chr XII) in the well was measured. On day 0 we assume that all of the *NTS2* signal in the well is associated with replicating Chr XII (rust, blue and green bars). The contribution of replicating ERCS (yellow bars) to the well-retained fraction in the gel was estimated by comparison with well-retained 2-micron on each day (B and D). The remainder of the NTS2 signal in the well (purple bars) is unaccounted for by either replicating Chr XII or replicating ERCs.

To determine whether the remainder of the rDNA copies were the result of replicating ERCs we again turned to the analysis of the 2-micron plasmids. One major difference between 2-micron and ERC replication is the fate of the two forks on the two plasmids. Each copy of the 2-micron plasmid is replicated once each cell cycle, like chromosomal DNA, (Zakian et al., 1979) by two bidirectionally oriented forks (Brewer and Fangman, 1987) while an ERC would be replicated by a single unidirectional fork (Brewer and Fangman 1988; Linskins and Huberman 1988). Thus, matched for size, an ERC would take roughly twice as long to complete replication as a 2-micron plasmid. We measured the signal for the *FLP1* probe in migrating forms of 2-micron and compared it to the signal retained in the well for each strain on each day (Fig 7B,D) and then doubled that value to estimate the proportion of ERCs that could be trapped in the well (yellow bars in Fig 7A,C). The sum of replicating Chr XII rDNA and replicating ERCs does not account completely for the total copies of *NTS2* in the well in aging cells, leaving the identity of a sizable fraction of *NTS2* signal unexplained (purple bars in Fig 7A,C). The fold change in *NTS2* signal amounts to >1100 copies of rDNA per cell for the *sir2Δ* and wt populations and >350 copies of rDNA per cell for the *fob1Δ* population.

### Assessing the cause of death in aging populations

To determine how the additional copies of the rDNA were distributed across individual cells and to determine whether the extra copies contributed to cell death we used flow cytometry and the non-specific DNA stain Sytox green to assess cellular DNA content across individual cells throughout the life span of our populations. We generated cell cycle profiles on each day for each of the three populations (Fig 8A; S4A Fig). These profiles consist of two main peaks, corresponding to the cells in G1 and G2 phases, with S phase cells in between. The day 0 samples are enriched for G1 cells (and are depleted for S-phase cells). In the following days, there was a return of cells with intermediate amounts of DNA between the G1 and G2 peaks, but most notably, in the *sir2Δ* mutant and to a lesser extent in wild type cells, subsets of cells were observed to contain more than two (and up to eight) genome equivalents of DNA (Fig 8A; S4A Fig). We quantified the proportion of the cells that contain more than 2 genome equivalents of DNA and found that more than 30% of the two *sir2Δ* population fell into this category by the end of life, and roughly 20% and 6% in wild type and *fob1Δ*, respectively (Fig 8B). Because some cells in the G2 peak may actually be G1-arrested cells with amplified rDNA, we may be underestimating the fraction of cells that have experienced an amplification event. The subset of cells containing more than 2C genomic content has an average genome size from 3.3 to 3.8 C, an increase in DNA content per cell that is consistent with the level of excess rDNA repeats estimated in Fig 6, top.

**Figure 8.**
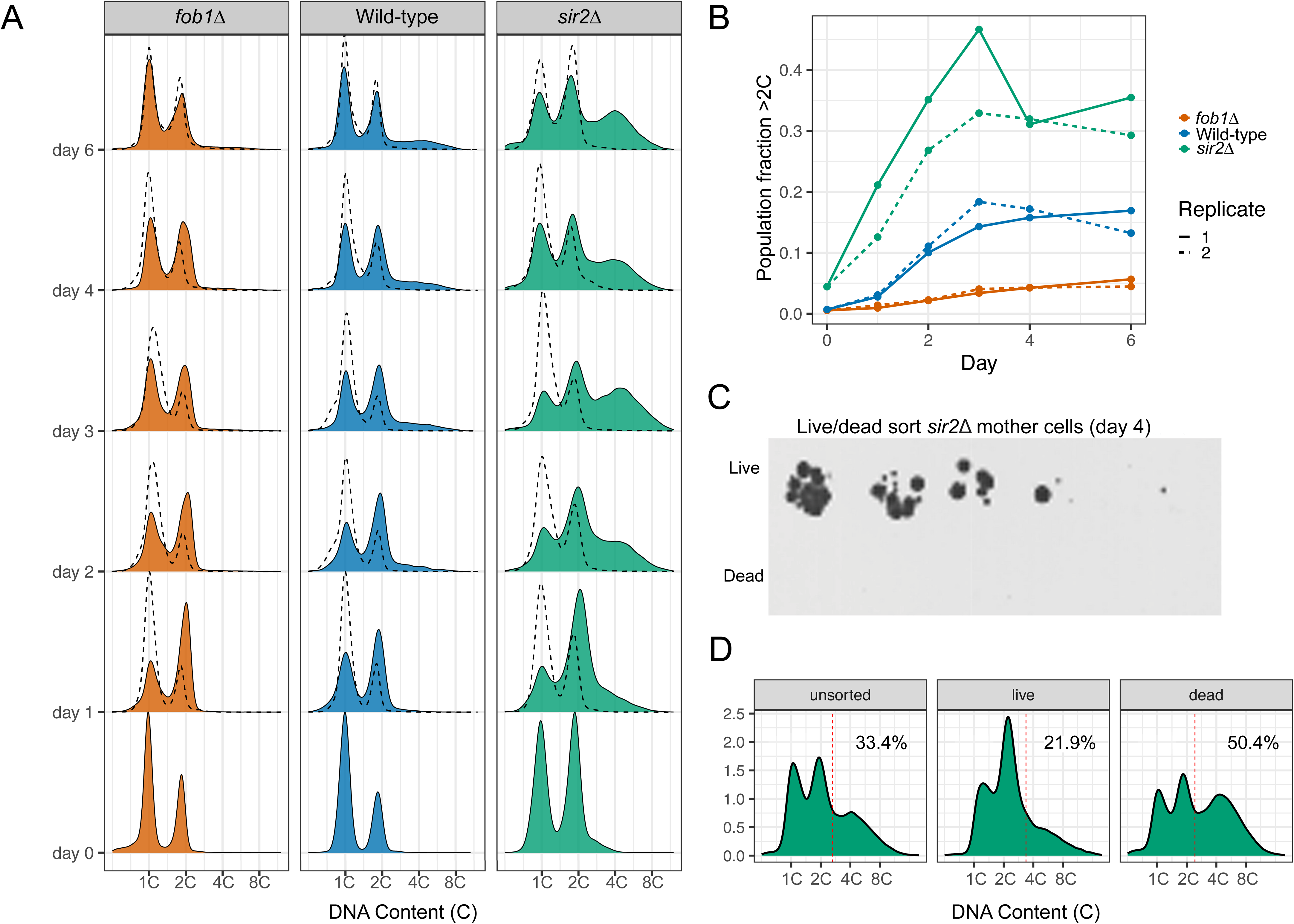
rDNA accumulation is heterogenous across cells from aging populations. A) Cell cycle distribution across the replicative lifespan of each population was generated by staining fixed cells with Sytox green and measuring fluorescence intensity with a flow cytometer (n > 10,000 cells). In addition to aging mother cells, we also measured the cell cycle profiles of daughters of aging mothers (dashed distribution). Day 0 cells were predominantly young mothers with an average of ∼1 bud scar per cell (Fig 3C). The x-axis was transformed from arbitrary fluorescence units to genomic content (C). The 1C and 2C peaks, corresponding to the genome before and after replication are present in each plot. In mother cells, only a subset of cells appears to accumulate additional DNA. Daughter cell cycle profiles (dashed lines) do not show evidence of DNA accumulation. B) The fraction of the population from the profiles shown in A (replicate 1) and in S4A Fig (replicate 2) with DNA content above 2.3N was quantified. C) Mother cells from the *sir2Δ* population were collected on day 4 and sorted into live and dead pools using fluorescence activated cell sorting (FACS) using the membrane permeable stain, propidium iodide. A spot dilution assay was used to validate the effectiveness of the sort. D) Cell cycle profiles were generated from unsorted, live and dead cells. Data for replicate 2 are shown in S4B Fig.

To determine whether the accumulation of additional rDNA was associated with a loss of viability, we used fluorescence-activated cell sorting (FACS) to separate the *sir2Δ* population into live cells and dead cells. The high efficiency of the sort can be seen in the spot dilution series shown in Fig 8C. We then generated cell cycle profiles for the live and dead populations and found that the non-viable population was more likely to contain additional DNA (Fig 8D; S4B Fig). This finding is consistent with rDNA accumulation reducing life span. Among the nonviable *sir2Δ* cells ∼50% have died without accumulating additional DNA. Could this population of cells be dying from a different form of rDNA instability occurring on Chr XII and is it causing Chr XII to be retained in the well to a greater degree than Chr IV (Fig 4E; S2E Fig)?

### Non-migrating rDNA species

DNA molecules can be retained in the well of a CHEF gel for a variety of reasons.

If incomplete lysis were responsible for the retention of Chr XII in the well, then we would expect the other chromosomes to show the same pattern, but they do not (Fig 4E, S2E Fig). A second possibility is that the non-mobile rDNA copies are due to the presence of ERCs larger than five repeats. However, we would predict that the larger an ERC is, the more likely it would be broken and produce a linear molecule upon death. We do not see signal in the range of discrete linear molecules that would be indicative of broken ERCs larger than five repeats (Fig 5C,D). A third possibility is that the well contains ERCs in the process of replication; however, our estimates for replicating ERCs, based on the proportion of replicating 2-micron (Fig 7B,D), accounts for less than half of the *NTS2* signal in the well by the end of their life span (Fig7A,C).

These considerations suggest that unexplained *NTS2* signal in the well is due to some other branched form(s) involving rDNA repeats. While unresolved recombination intermediates could contribute to the *CDC45* probe in the well, they do not provide an explanation for the massive increase in the *NTS2* signal in the well. Is there a way to account for 1) the increase in proportion of Chr XII in the well; 2) the large increase in unexplained rDNA repeats in the well; and 3) the death of cells without an apparent increase in DNA content? We hypothesize that unresolved replication forks within the rDNA could be the cause—specifically, a replication fork that does not have an oppositely oriented fork for termination. An un-terminated replication fork in the rDNA could explain all three features, but what event could leave behind an unopposed replication fork?

### Models to explain the “General Theory of rDNA Aging”

In 1986 Bruce Futcher proposed a model that explains the amplification of the 2- micron plasmid. It involves site-specific recombination between inverted repeats that differ in replication status (S5 Fig). This recombination event converts the two bidirectionally orientated replication forks into two forks that are moving in the same direction around the 2-micron circle. Instead of terminating at a site opposite the origin, the two forks are unopposed and continue copying the plasmid multiple times without requiring a new round of replication initiation. Could something similar be occurring during repair of broken forks in the rDNA—repair of a broken fork using a repeat that generates unopposed replication forks? And could these aberrant repair events lead to retention of Chr XII in the well, the unexplained additional rDNA repeats, and the death of cells without a massive increase in rDNA copies?

### 1. Catastrophic IntraChromatid Repair (CICR)

In Fig 2 we considered outcomes for replication fork restoration using an un- replicated nearby repeat. However, because replication initiation events in the rDNA are scattered across the entire locus with approximately one in five of the origins being active and the other nearby origins being passively replicated (Fig 1; Brewer and Fangman, 1988; Linskens and Huberman, 1988), there are many opportunities for a broken fork to use a replicated repeat for repair instead. We explored the possible consequences of such a repair (Fig. 9)—i.e. the outcome of the Futcher model when applied to direct, chromosomal rDNA repeats.

**Figure 9:**
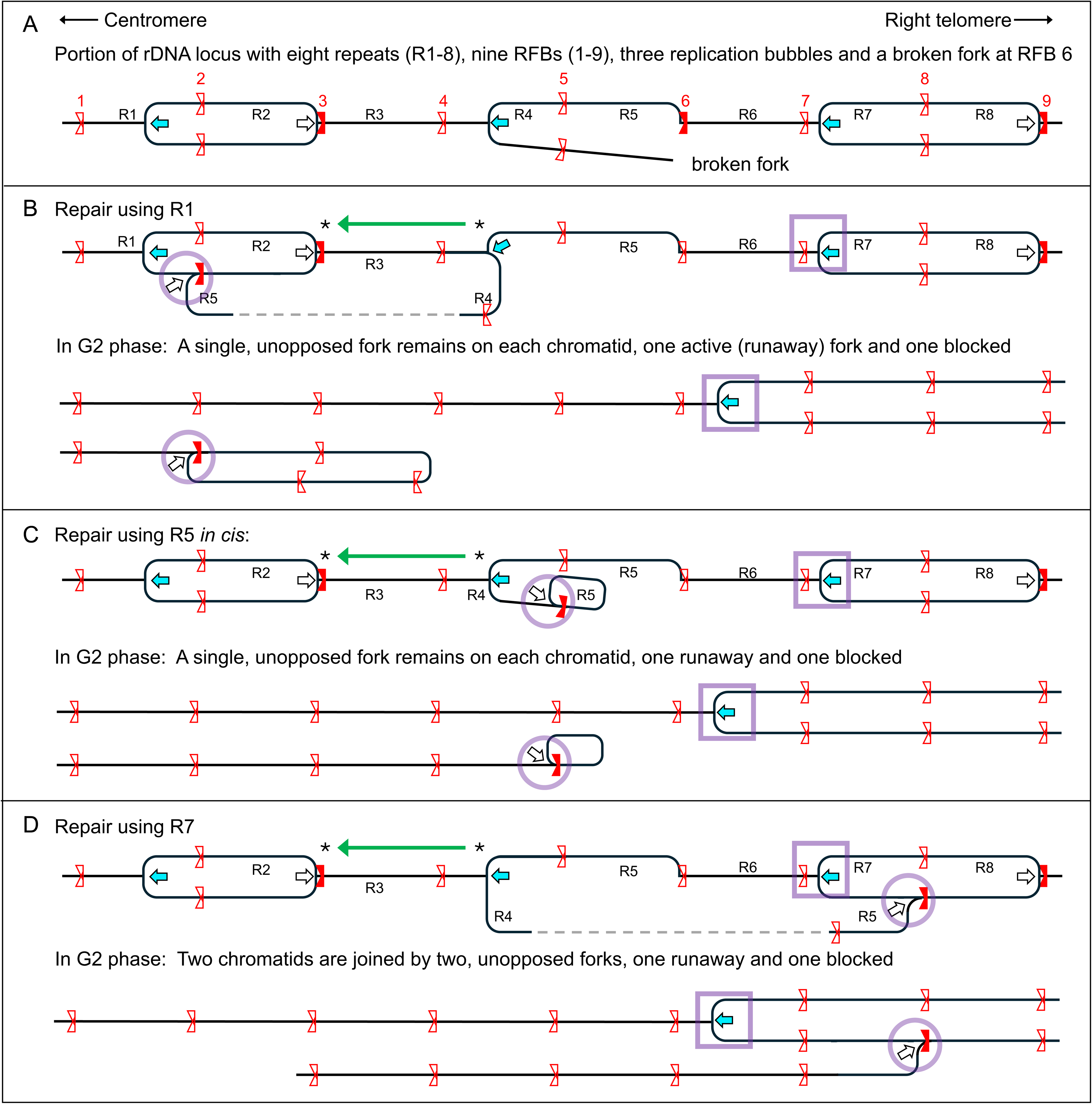
Catastrophic IntraChromatid Repair of a broken replication fork (CICR) A) A section of the rDNA with eight repeats (R1-R8), three replication bubbles, and a broken fork at RFB6 is illustrated. B) Replication fork restart using R1 as the repair template is illustrated. The active fork in R4 will replicate through R3 (green arrow) and terminate with the arrested fork at RFB 3. The repair leaves two replication forks un-paired (blocked fork, violet circle; runaway fork; violet square). After all other bidirectionally oriented forks have terminated on Chr XII the two chromatids of Chr XII will have single remaining forks that are unpaired. All sequences distal to the rDNA are unevenly distributed between the two sister chromatids: a truncated Chr XII that ends in a lariat in the rDNA and the other chromatid with two distal segments joined by a runaway fork within the rDNA locus. C) Fork repair using R4 *in cis* produces the same two types of chromatids (as produced by in B) after completion of all bidirectionally oriented forks. D) Repair using a distal repeat (R7) also produces unpaired blocked and runaway forks but in this case after completion of all other forks, the two chromatids remain tethered by a linker of rDNA repeats that will expand in length as the runaway fork continues replication. These two sister chromatids would generate an anaphase bridge during mitosis.

There are two possible orientations for interactions between a broken fork and a replicated rDNA repeat, use of either a centromere-proximal or centromere-distal replicated repeat as the repair template (Fig 9A). First, consider the use of a centromere-proximal, replicated repeat as the repair template (Fig 9B). Restoration of the fork and termination of all paired, bidirectionally oriented forks along Chr XII results in complete replication of Chr XII with two-unpaired forks remaining. The sequences are not distributed equally to the two chromatids: one chromatid contains all sequences to the left of the rDNA but terminates in a loop of rDNA repeats with the branch-point at an RFB-arrested fork; it is missing all sequences to the right of the rDNA. The other chromatid runs the entire length of Chr XII but all sequences distal to the rDNA are duplicated, attached by a replication fork in the rDNA and share a common centromere. This fork, in the permissive orientation with respect to the RFB, can continue synthesis toward the centromere in an unchecked manner as a “runaway fork”. Because of their branched structures (lariat and Y-shaped chromatids) we anticipate that they would be retained in the CHEF gel well. In addition, the runaway fork on the Y-shaped chromatid could generate an increase in rDNA repeats that are not ERC-related. Note, that the same structures result if the broken fork is repaired *in cis* using an adjacent repeat on the fragmented arm of the replication bubble (Fig 9C). We refer to these events—involving RFR between the broken fork and an adjacent replicated or *in cis* repeat—as CICR events, for Catastrophic IntraChromatid Repair.

The second possible interaction in a CICR event can be between the broken fork and a distal replicated repeat (Fig 9D). After completion of all opposed replication forks, this repair produces chromatids that remain connected by two diverging replicating forks, one arrested at an RFB and the other creating a runaway fork proceeding towards the centromere. Because of its branched structure these unresolved chromatids would remain in the well of the CHEF gel and the runaway fork could be contributing extra rDNA repeats to the well.

When cells with either CICR event attempt mitosis, we predict that the results would be catastrophic for these haploid cells. In the first two examples one of the cells would inherit a truncated Chr XII missing unique genes distal to the rDNA and would be lethal (Fig 9B,C). The other cell would inherit a branched Chr XII with duplicated copies of the distal unique genes. This branched chromosome would be self-perpetuating in subsequent cell cycles leading to continuous aneuploid state, assuming the runaway fork fails to trigger a checkpoint to prevent cell division. A CICR event that involves repair from a distal replicated repeat (third example; Fig 9D) produces attached chromatids that would generate an anaphase bridge and both cells receiving broken chromatids would have to deal with a double stranded break and single unopposed replication forks.

### 2. Replication Fork Repair involving ERCs

ERC replication is expected to be unidirectional with one fork that is moving around the circle and the other fork that is blocked at the RFB. If the stalled fork is broken, the remaining fork can continue around the circle in a form of rolling-circle replication (S6A Fig) with the un-protected broken end being a target for exonucleases. However, if the break is repaired using a repeat on the ERC (adjacent to RFBs 1, 2, or 3) then a variety of outcomes are possible. These three choices restore the balance of oppositely oriented forks and may or may not result in changes in the size of one of the replicated ERCs (S6A-C Fig).

There is one additional, intramolecular repair event involving an ERC with a broken fork (Fig 10). If the repeat *in cis* to the broken end (adjacent to RFB4) is used for repair then the two forks are oriented away from one another generating a linear molecule of just rDNA repeats with lariats at both ends (a double-lariat ERC or DL- ERC). One fork is arrested at the RFB and the other is a runaway fork in a rolling-circle configuration (Fig 10A). This molecule would also likely be retained in the well and be retained by the mother cell at mitosis. Continued replication from the runaway fork would add to the length of the linker between the two repeats (Fig 10B). In subsequent cell cycles, new initiation events could occur in either lariat or in the linker between them. Initiation events in either lariat preserve the lariat structure but add repeats to the linker. However, an initiation event in the linker creates a unique structure: it generates a new pair of blocked and runaway forks (Fig 10C) nested between the original two divergently-oriented, unpaired forks. Each cell cycle, the probability of initiation within the linker increases because of its ever-increasing length and the number of runaway forks expands exponentially from that initial repair event (Fig 10A). (Note, DL-ERCs can also be produced when the broken fork is repaired using a second ERC (Fig 10D)). These runaway forks are reminiscent of the forks in the Futcher model of 2-micron amplification (S5 FIG) where they continually chase each other around the circle, but in the case of the DL-ERCs, they chase each other towards the lariat which always remains just out of reach.

**Figure 10:**
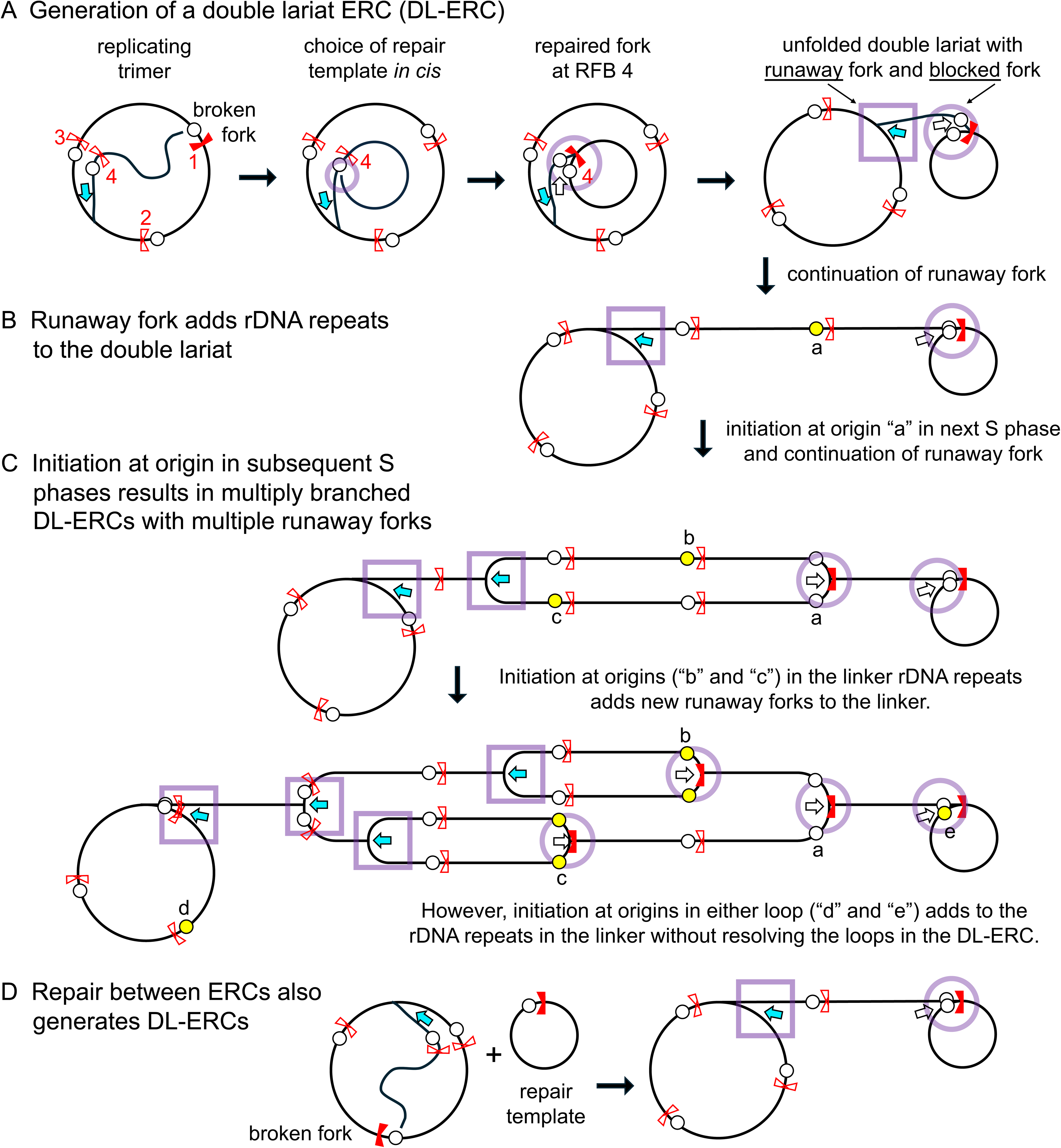
Repair of a broken fork in an ERC results in a Double-Lariat ERC (DL- ERC). A) A replicating ERC with three repeats (trimer) suffers a broken fork at RFB1 that can be repaired using a proximal replicated repeat *in cis* (adjacent to RFB4) as the template. The two forks, one blocked at RFB4 and the other continuing around the ERC, will no longer meet for termination. Unfolding the nested circles reveals a double-lariat (DL) ERC. Active origins are shown as yellow circles; potential origins are shown as open circles. B) The one active fork on the DL-ERC can continue to add repeats to the linker by circumnavigating the trimer. In the following cell cycles the origins are re-licensed and any of them (ie, “a”) can initiate replication. C) Activation of origin “a” generates two forks in the linker with neither of them being opposed by a fork in the opposite direction. Meanwhile, the runaway fork produced in B continues to add repeats to the linker. In later cell cycles, additional origins in the linker (“b” or “c”) add additional runaway and blocked forks to the DL-ERC resulting in a multiply branched linker with lariats at each end. Initiation in either loop (“d” or “e”) does not resolve the loop but only adds additional repeats to the linker. D) Repair of the broken fork of an ERC can use a second ERC for repair and generate the same DL-ERC as shown in (A).

Finally, an ERC with a broken fork could also use a chromosomal repeat for repair (S7 Fig). The use of an unreplicated repeat as the repair template (S7A Fig) generates unbalanced chromatids, similar to the ones described in Fig 9B,C, but with the opposite polarity: the truncated partial chromatid, ending in a loop in the rDNA, does not contain a centromere and the Y-shaped chromatid has two centromeres.

Attempts to segregate these two Chr XII “chromatids” at mitosis would be catastrophic. The use of a replicated repeat as the repair template (S7B Fig) produces two complete Chr XII chromatids with one of them containing a side branch with two unresolved forks. Like the DL-ERCs, this tethered rDNA lariat could grow in length and complexity due to additional origin initiation events within the tether. Either of these repair events would result in a chromosome that is retained in the well and a runaway fork that could continue to generate rDNA copies long after S phase is completed.

## DISCUSSION

Aging and rDNA instability have largely been connected through the ERC hypothesis. Mother cells form ERCs, accumulate them, and eventually succumb to their toxic effects. ERCs can be thought of as a chronic problem needing multiple generations to become symptomatic. In our models, we propose that a broken fork in the rDNA is capable of generating a variety of rDNA rearrangements in addition to ERCs which may shorten life span in different ways—by the sheer number of extra rDNA copies or by generating branched forms of Chr XII that are lethal during mitosis.

The key feature of these models is the choice of the repair template. Using a repair template *in cis* or one that differs in replication status generates unopposed replication forks with unique consequences.

The completion of chromosome replication requires that each fork established from an origin of replication either meets a fork in the opposite direction for termination or runs off the nearest telomere. An unfortunate choice of repair template can disrupt this delicate balance. CICR events involving the chromosomal rDNA locus create chromosome imbalances that would be acutely lethal in haploids. While contributing transiently to an excess of rDNA repeats in the well, they can explain the excessive retention of Chr XII in the well and the death of cells without substantial rDNA accumulation. DL-ERC events, like the Futcher model, generate unopposed runaway forks that chase one another along the template toward, but never catching up to, the terminal lariat. DL-ERCs can explain the excessive, unexplained *NTS2* hybridization we observe in the well (Fig 7A,C) and why these molecules are not released as discrete linear fragments after cell death (Fig 4A,D and S2A Fig): a single double-stranded break cannot convert DL-ERCs into discrete linear forms that are able to enter the gel.

In this work we have presented our models from the perspective of repairing the break of an RFB-arrested replication fork; however, any fork, arrested or not and even moving in the permissive direction in the rDNA, could undergo similar CICR or DL- ERC events with similar outcomes. In addition, a double-ended, double-stranded break repaired as a double-Holliday junction can generate unopposed replication forks if the repair template is in a different replication state. We chose to illustrate only repair of the fork arrested at the RFB because it simplifies the diagrams.

### Roles for Sir2 and Fob1 in the generation of runaway forks

Replication fork breakage and repair is stochastic in nature, but the frequency of breaks and/or the choice of repair template are influenced by the *SIR2* and *FOB1* genotypes. The effect of genotype on the frequency of rDNA instability is evident in our analysis of genome content by flow cytometry (Fig 8; S4 Fig), in the rate that ERCs are generated (Fig 5A; S3A Fig; Fig 6), and in the accumulation of excess rDNA repeats in the CHEF gel wells (Fig 6). The types of events do not differ among the three strains, but the rate of occurrence across the different genotypes does. How might the absence of Fob1 and Sir2 fit with our mechanistic models for rDNA instability?

Fob1 binds to sequences downstream of the 35S transcription unit and through its binding at the RFB prevents passage of the replication fork that would meet RNA Pol1 head-on. As a consequence the fork persists at that site until it is terminated by a fork moving from an adjacent replicon. This persistence is thought to make the fork fragile and in need of repair. This RFB region also contains the *HOT1* locus that stimulates recombination between adjacent repeats (∼150-fold when Fob1 is present; Ward 2000). However, is it unclear whether Fob1’s role in fork breakage and recombination stimulation are one and the same: stimulation of recombination between flanking sequences is equivalent in the fork blocking direction and the permissive orientation with respect to the adjacent replication origin (Ward 2000) arguing that a stalled fork, and its subsequent breakage, is not necessary for the enhanced recombination activity of *HOT1*. Fob1 has also been shown to dimerize and the role of Fob1 in recombination enhancement might be in its role of bringing two repeats together (Choudhury et al., 2015). Other possible roles of Fob1 could be in recruiting other proteins that break branched DNA or in interfering with subsequent repair.

Regardless of the mechanism, in a *fob1Δ* strain chromosomal rDNA copy number variation, as well as DL-ERCs and CICR events still occur, but a reduced frequency.

Sir2 is a histone deacetylase first discovered for its role in mating type silencing Klar et al. 1979; Rine et al. 1979). Subsequently it was shown to repress PolII transcription in the *NTS1* region of the rDNA repeats (Kobayashi & Ganley 2005). In the absence of Sir2 the time of replication of the rDNA locus is advanced with more origins firing early in S phase (Pasero et al. 2002; Kwan et al., 2023) and cohesion is no longer able to enforce the registration between sister chromatids in the rDNA locus (Kobayashi & Ganley 2005). This combination of phenotypes creates a situation where the choice of repeat used for repair is broadened and the broken fork has a greater possibility of choosing a replicated repeat as the repair template. For example, *sir2Δ* cells produce fewer monomer ERCs relative to WT (Fig 5A and S3A Fig) consistent with the suggestion that in *sir2Δ* cells a broken fork in the rDNA has greater mobility to engage a more distant repair template when forming an ERC. Also, the well-retained rDNA copies appear sooner in the *sir2Δ* strain than they do in wild type cells (Fig 6), perhaps because of the shift to earlier and increased initiation events at origin in the rDNA repeats (Pasero et al. 2002; Kwan et al. 2023).

### Observations from the literature

We are not the first to analyze genomic DNA from aging populations of yeast cells on CHEF gels, nor are we the first to generate hypotheses on what is “stuck in the well.” Previous analyses of non-migrating DNA have treated it as a measure of branched recombination intermediates and quantified the rDNA signal retained in the well as an index of rDNA instability (Ganley et al., 2009). Others have found that compared to a probe for chromosome II, chromosome XII was four times more likely to be retained in the well and also attributed this increase in non-migrating DNA to be the result of rDNA recombination intermediates (Pal et al., 2018). Our quantification and kinetic analysis of rDNA forms suggest the well-retained rDNA is a mixture of at least four species—normal replicating chr XII and unidirectionally replicating ERCs, but also CICR events and DL-ERCs that both contain runaway forks, in addition to Holliday junctions.

Aging experiments on additional mutants are also consistent with our models of rDNA instability. First, in a strain with impaired rDNA origins (*rARS-Δ3*; Hattori et al., 2022) ERC production in mothers (∼7 divisions) is significantly reduced yet the accumulation of additional rDNA species in the well of the CHEF gel is comparable to that of wild type cells. This finding is consistent with the proposal that the well-retained rDNA in the *rARS-Δ3* strain is not in the form of autonomously replicating ERCs but some other form of branched rDNA such as those produced by a CICR event.

Second, the *rtt109Δ* mutation significantly shortens RLS (Feser et al., 2010) and increases rDNA instability (Ide et al., 2013). Using atomic force microscopy to examine DNA that remained in the well of a CHEF gel from the *rtt109Δ* mutant, Ide et al. (2013) found lariat structures, with a loop size equal to roughly one rDNA repeat, at a frequency of ∼1% in the *rtt109* mutant and at a frequency of <0.1% in wild type cells.

They proposed the lariats arose as follows: “To initiate rolling circle replication, the broken end at the RFB must recombine with . . . popped-out circular molecules (ERCs).” However, the fork established by this invasion event would be in the non-permissive orientation for replication with respect to the adjacent RFB and we believe would be unlikely to form a rolling circle replicon. We suggest that their lariats could have been generated by CICR events or DL-ERCs.

Third, Zylstra et al. (2023) measured the DNA content of *rad52Δ* mutant diploid cells as they aged and found no ERCs, suggesting that Rad52 is required for ERC formation. By whole genome sequencing, they found a modest increase in the unique region of Chr XII between the rDNA and the right telomere in *rad52Δ* mutants (S8A Fig). In addition, they observed that Chr XIIr also accumulated with age in wild-type cells (S8B Fig) and suggest that the amplification of Chr XIIr may be a general aging phenotype. They concluded that the extra sequences were due to a linear fragment of 1.8 Mb but they unfortunately did not include a CHEF gel to confirm their suspicions. It is possible that the overabundance of Chr XIIr was the consequence of a CICR event, but it remains unclear what role *rad52Δ* might play in CICR events. The Tyler lab also reported an increase in Chr XIIr in aging haploids (Hu et al., 2013) but attributed the increase in read depth to translocations between the rDNA and other sites in the yeast genome.

To determine whether the aged populations in our study also amplified Chr XIIr, we sequenced the entirety of our time course for replicate 1. We found no indication of an amplification of Chr XIIr (S8C-E Fig). We did not see similar amplification in Chr XIIr in our aging populations perhaps because the cells were haploid and CICR events would be acutely lethal. In contrast the populations sequenced by Zlystra et al. (2023) were diploids which are more tolerant of chromosome aneuploidies and aneusomies.

Finally, there is also precedence in the literature for the basic feature of the CICR model that intramolecular recombination between repeats of different replication states alters the orientation of replication forks. Bruce Futcher model for amplification of the yeast 2-micron plasmid (1986; S5 Fig) provided the inspiration for us to ask what the outcome of recombination between direct repeats of different replication status would have on chromosome structure. While the recombination event in 2-micron is site specific and occurs between inverted repeats, the resolution of this cross-over event converts the bidirectionally moving replication forks into two rolling circle forks that chase each other around the plasmid spinning out multiple tandem copies of the 6.3 kb repeat. A second model by Johansson and Diffley (2021) proposes a scenario involving DNA repair and replication fork restart that can result in DNA re-replication. They did not specifically address issues at the rDNA, nor address the consequences on chromosome segregation. However, when we apply their model to the rDNA locus, we predict that it would generate a product identical to the CICR event described in Fig 9D; chromatids that are held together by a tether that ends in one blocked fork and one runaway fork, unopposed and oriented away from each other. The key similarities shared by this model, in which a replicated segment recombines with a re-replicated segment, and the Futcher and CICR models, are the creation of unopposed replication forks and the consequences that this imbalance has on genome stability.

### Consequences beyond the rDNA locus

Because we were investigating yeast aging, we focused our attention on repair of a broken fork that occurs in the rDNA and propose that the choice of repair template (replicated vs. unreplicated or *in cis* to the break) is crucial. However, the repair processes we have described could hypothetically occur between any sequences that are repeated and that differ in replication status at the moment of the repair. We suggest that fork repair using dispersed repeats (such as Ty elements in yeast) could not only generate translocation (which are not necessarily lethal until cells undergo meiosis) but more acutely, could generate the same kind of branched or truncated chromatids with unopposed replication forks. Broken forks could use dispersed repeats on the same or different chromosome for repair and even breaks in non-repeated regions could use regions on the homologue if the two regions are asynchronously replicated.

Unfortunately, a discussion of these potential events and the possible safeguards that cells have evolved to prevent them is beyond the scope of this manuscript.

## METHODS

### Strains used in this study

^Wild-type (WT)^ FY4 - *MATα*, *flo1*::KanMx, *flo9*::NatMx

*sir2*Δ FY4 - *MATα*, *flo1*::KanMx, *flo9*::NatMx, *sir2*::KanMx

*fob1*Δ FY4 - *MATα*, *flo1*::KanMx, *flo9*::NatMx, *fob1*::KanMx

Each of the strains used in this study contains deletion of two genes which contribute to flocculation. The deletions of *FLO1* and *FLO9* facilitate the separation of mother cells from daughter cells. Although the laboratory strain we used, FY4, is not known to flocculate in standard synthetic growth media, flocculation can be induced in laboratory strains with age (Hendrickson et al., 2018). We also attempted to keep rDNA copy number consistent between the three strains by selecting transformants with similar rDNA copy number to our WT strain. However, because of the inherent rDNA copy number variation in *sir2Δ* strains, we were only partially successful. By the time we analyzed the cells on day 0, Chr XIIs was already a heterogeneous mixture of sizes with a mean rDNA copy number in the two replicates of ∼90 copies.

### Magnetic labeling and cultivating aging yeast

Cultures derived from single cells were grown overnight in 2 mL synthetic complete (SC) media containing 2% glucose. The overnight cultures were transferred into 25 mL SC media and allowed to grow overnight until saturation. For each strain and each replicate, 10^9^ cells (∼50 ODs) were isolated and washed five times in 5 mL phosphate buffered saline (PBS) before being resuspended in 5 mL PBS + 30% polyethylene glycol (PEG; w:v). To activate the carboxylate modified magnetic beads, 1-Ethyl-3-(3-dimethylaminopropyl)carbodiimide (0.091 g) and N-Hydroxyl succinimide (0.541 g) were added to 2 mL MES buffer (1X) along with carboxylate modified magnetic particles (Seramag speedbeads, 0.2 mL, washed twice in 0.5 mL MES). The mixture was briefly sonicated before it was placed on an orbital shaker at room temperature for 45 minutes. After the incubation, the beads were washed three times in 1 mL PBS (1x) and sonicated again before they were added to the yeast cells suspended in PBS+30% PEG. The cells and beads mixtures were then placed on an orbital shaker at room temperature for 45 minutes. The magnetically labeled cells were isolated with a magnetic rack and washed five times in 25 mL SC media before being resuspended in 1mL SC media. A small aliquot (50 μL) of these cells was saved for characterization and the remaining portion was loaded into two chemostats. The labeled cells were allowed to adhere to the exterior magnets for 30 minutes before the media and air pumps were turned on. SC medium was perfused into the cultures at a rate of ∼33 mL hr^-1^ which corresponds to a dilution rate of ∼2 hr^-1^. Filtered air was continuously perfused into the chemostats to maintain positive pressure inside the vessel.

### Mother cell sampling and purification

The aging populations were sampled over a six day period by first stopping the media and air pumps and agitating the chemostats to free the cells from the walls of the chemostat. At each sample time ∼10^7^ cells were isolated. The mother cells were separated from the daughter cells using a benchtop magnet. Mother cells were washed on the magnet five times with 1 mL PBS.

### Viability and bud scar quantification

Viability was approximated using the membrane permeability stain propidium iodide. Briefly, 5 μL of propidium iodide (1 mg/mL) was added to 1 mL PBS containing 5x10^5^ cells. Cells were left in the dark for 20 minutes before viability was assessed using a C6 flow cytometer. For each sample >20,000 cells were measured.

Cells were first gated from the free magnetic particles and viability was assessed using the FL3 channel (630 nm).

The average replicative age of the culture was assessed using wheat germ agglutinin conjugated to alexafluor 488 (WGA-488). 5x10^5^ fixed cells were washed in 1 mL PBS and sonicated before 5 μL 1 mg/mL WGA-488 was added and the culture was left to stain for 20 minutes. The cells were centrifuged and resuspended in 20 μL PBS. Z-stack images of the stained cells were collected using a Leica widefield microscope at 100x. The bud scars of 40 cells were quantified for each sample.

### Analysis of genomic DNA by CHEF gel electrophoresis and Southern blotting

Mother cells retrieved from the aging chemostats were embedded in 1% GTG low-melt agarose (10^7^ cells for day 0 and 5x10^6^ cells for the remaining days in a 90 μl plug). Genomic DNA was purified *in situ* according to Tsuchiyama et al. (2013). The plugs were embedded in a 1% LE agarose gel and electrophoresis was carried out in 2.3 L of 0.5 × Tris Borate EDTA (circulating at 14°C) using a Bio-Rad CHEF DR-II electrophoresis cell. Conditions for electrophoresis that maximized the resolution of Chr XII were 100 V for 68 hr, with a switch-time ramped from 300 to 900 sec (Kobayashi et al., 1998). The gels were stained with Ethidium Bromide (0.3 mg/ml) for 30 minutes, destained for 30 minutes and photographed with UV illumination. Southern blotting to GeneScreen membranes and hybridization with ^32^P-dATP labeled PCR or restriction fragment probes were carried out using standard protocols. We sequentially probed the blots with different labeled fragments after stripping twice in boiling 0.1xSSC, 1% SDS. The probes included fragments to unique sequences on Chr IV (*GAL3*), Chr XII (*CDC45*), Chr X (*ILM1*), Chr V (*CUP1*), and repeated sequences on 2-micron (*FLP1*) and the rDNA (*NTS2*). To image and quantify the signal intensities of individual bands and/or lanes, we exposed the blots to X-ray film and to Bio-Rad Molecular Imaging FX phosphor, respectively. Phosphor screens were scanned using a Bio-Rad Personal Molecular Imaging scanner and analyzed using Bio-Rad’s Quantity One software.

**Table 1.**
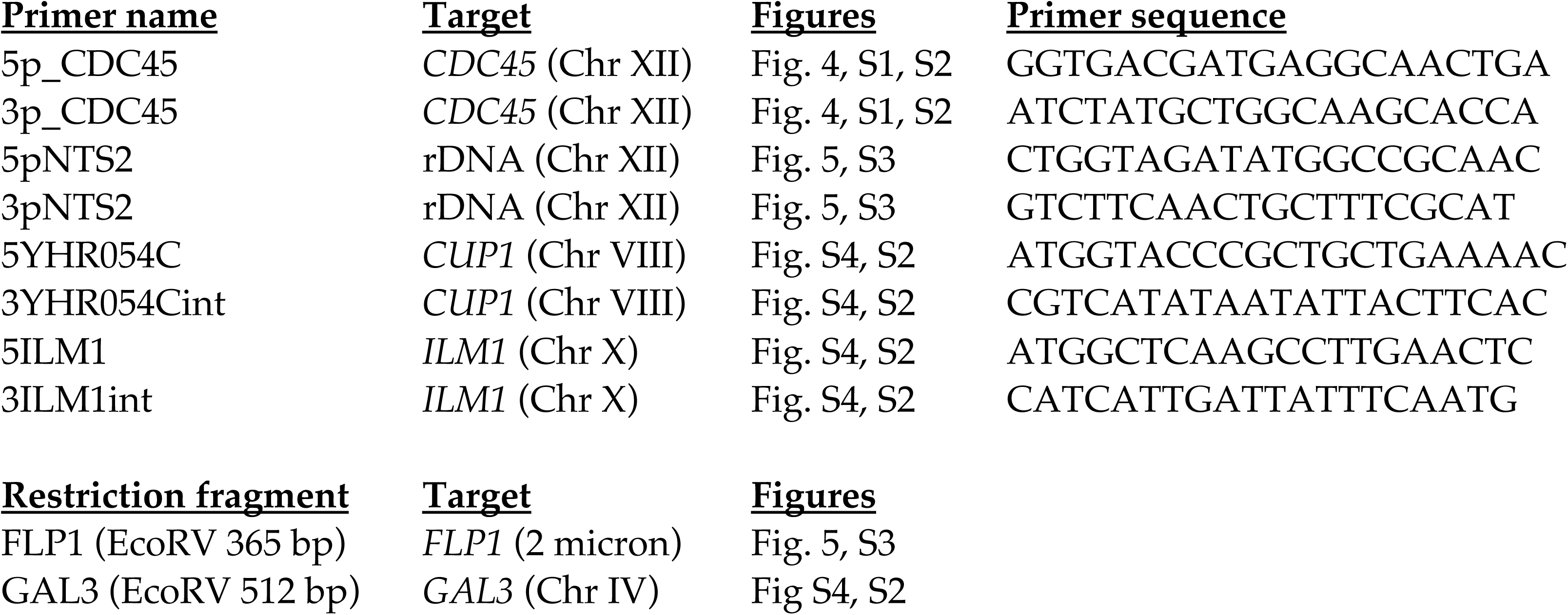
Southern blot probes.

### Live/dead sorting

To assess viability, 5 μL of propidium iodide (1 mg/mL) was added to 1 mL PBS containing 5x10^5^ purified mother *sir2*Δ cells and incubated in the dark for 20 minutes.

The population viability was assessed to be ∼40%. 100,000 cells assessed as dead by the propidium iodide uptake were collected along with a fraction of 100,000 cells assigned as live by the lack of fluorescence using a Sony MH800 fluorescence assisted cell sorter. A series of 10-fold dilutions was spotted onto YPD plates to assess the efficacy of the sort.

### Sytox green staining

At each sampling time ∼10^5^ cells were fixed in 70% ethanol. Cells were pelleted and washed twice in 1 mL 50 mM sodium citrate, treated with RNase A (0.25 mg/mL) and incubated at 50°C for 1 hour. Cells were then treated with proteinase K (0.5 mg/mL) and incubated at 50°C for an additional hour. Sytox green was added to the samples which were stored at 4°C until they were measured on a Becton Dickinson Accuri C6 flow cytometer.

### Illumina sequencing and copy number quantification

Libraries were prepared with the Illumina DNA Prep kit and sequenced as paired-end reads on a NextSeq 2000. Reads were aligned to the *S. cerevisiae* S288C reference genome (sacCer3/R64-1-1) with BWA-MEM. Mean read depth was computed in 5-kb windows across the Chr XII and normalized to the median window depth of the rest of the genome. Single copy coverage ranged from 34 to 200-X.

## ACKNOWLEDGMENTS

The authors would like to thank Lorraine Symington, Bruce Futcher and Jocelyn Verhey for helpful discussions, Josh Cuperus for reading an early version of the manuscript, Bruce Futcher for insightful comments on the first version of the manuscript, Rebecca Martin for help with Southern probes, and Kelsey Zane for copy editing of an early version of the manuscript.

## DATA AVAILABILITY

The raw data for Southern blot quantifications, flow cytometry, viability staining and bud scar counts can be found on FigShare (DOI : 10.6084/m9.figshare.34325157).

The raw whole genome sequencing data will be made available upon acceptance.

## SUPPLEMENTARY FIGURE LEGENDS

**Supplementary Figure 1:**
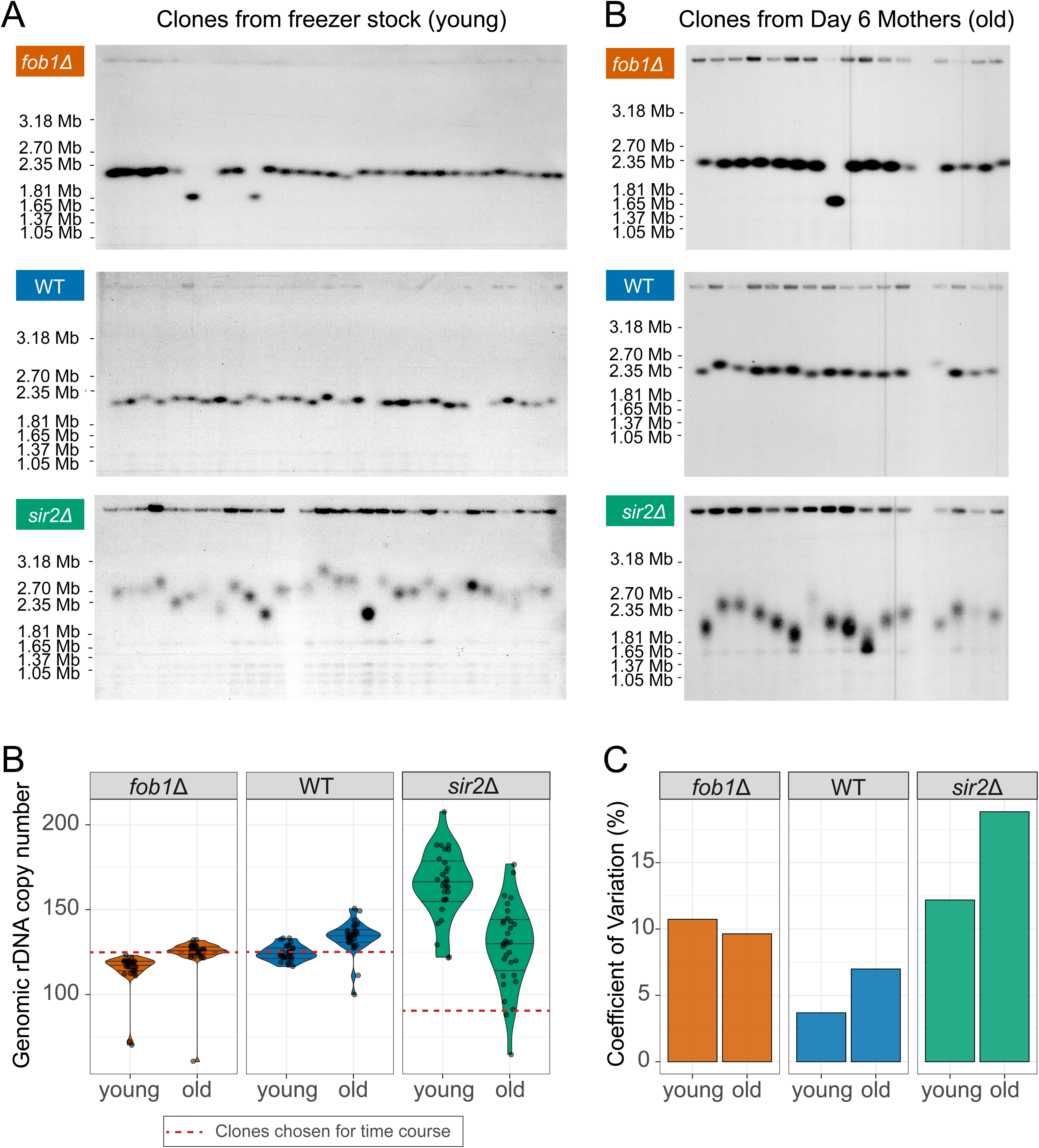
Variation in rDNA copy number among clonal isolates. A) 28 colonies derived from isogenic freezer stocks of *fob1Δ*, wild-type (WT) and *sir2Δ* cells were grown from single cells and karyotyped by CHEF gel electrophoresis. B) 16 colonies derived from day 6 (old) mother cells of the three strains were analyzed in a similar manner on CHEF gels. The Southern blots (A and B) of the CHEF gels were probed with *CDC45* to determine the size of Chr XII. C) From the size of Chr XII we estimated the rDNA copy number of each of the 28 colonies of each genotype shown in A. D) The coefficient of variation of chromosomal rDNA copy number increases in clones from old *sir2Δ* and WT mothers.

**Supplementary Figure 2:**
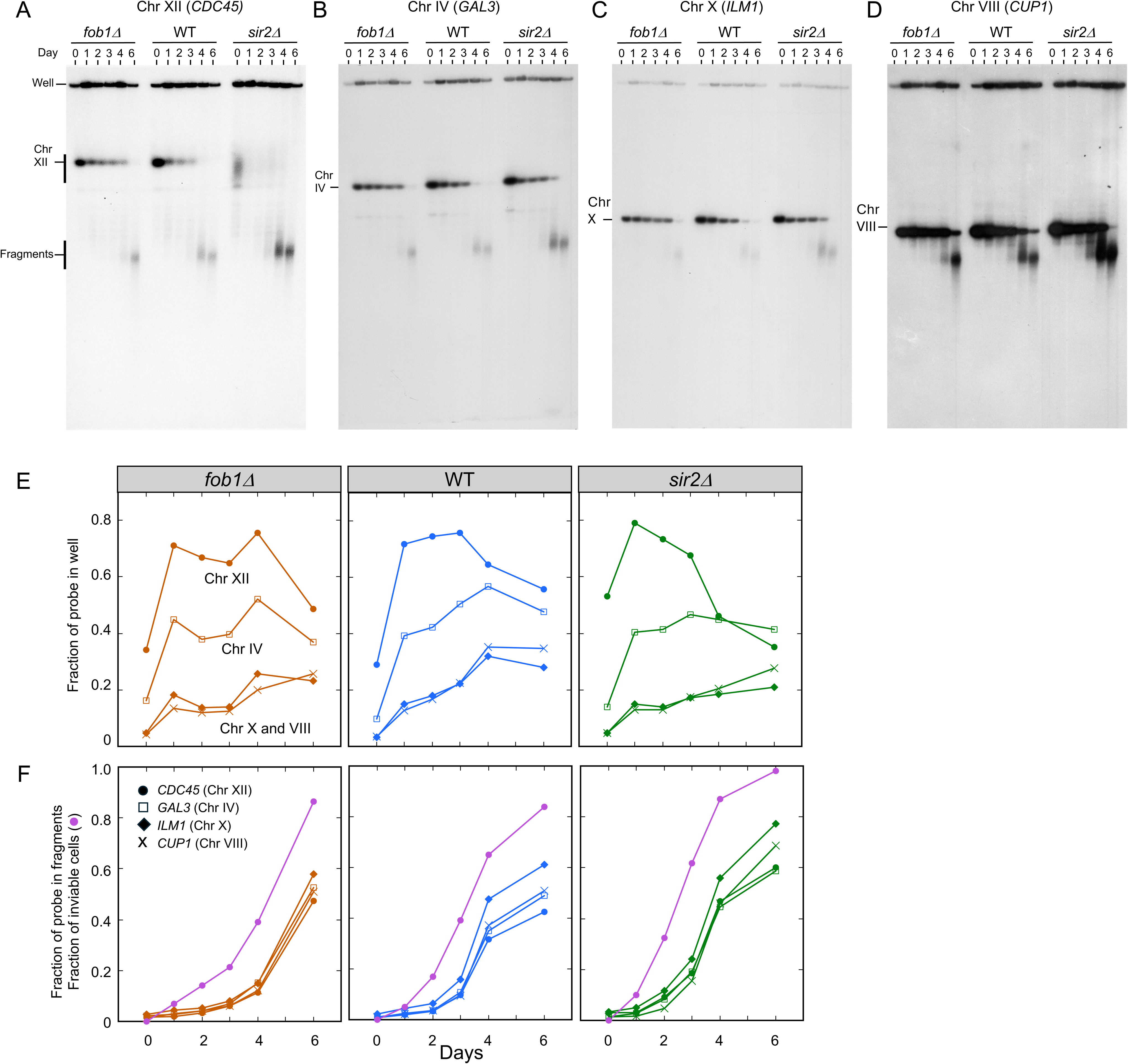
Characterizing individual chromosomes and their distribution in the CHEF gels of replicate 2. This figure is an identical characterization of replicate 2 of the aging chemostat populations using probes from four different chromosomes. Refer to the legend of Fig 4 for details.

**Supplementary Figure 3:**
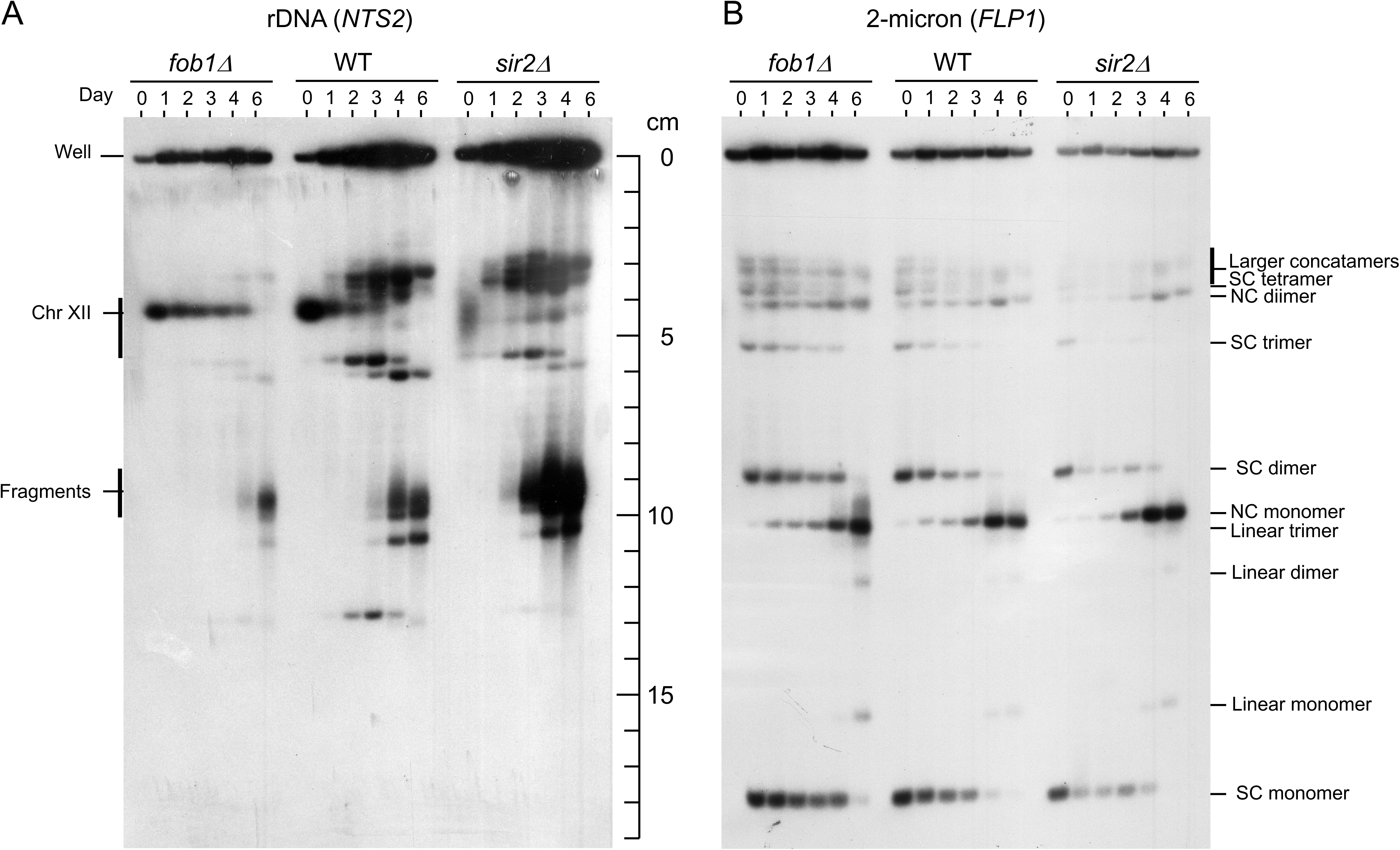
Southern blot analysis of replicate 2 aging populations. The Southern blot of CHEF gel 2 was hybridized with the *NTS2* and *FLP1* probes as described in Fig 5A and B.

**Supplementary Figure 4:**
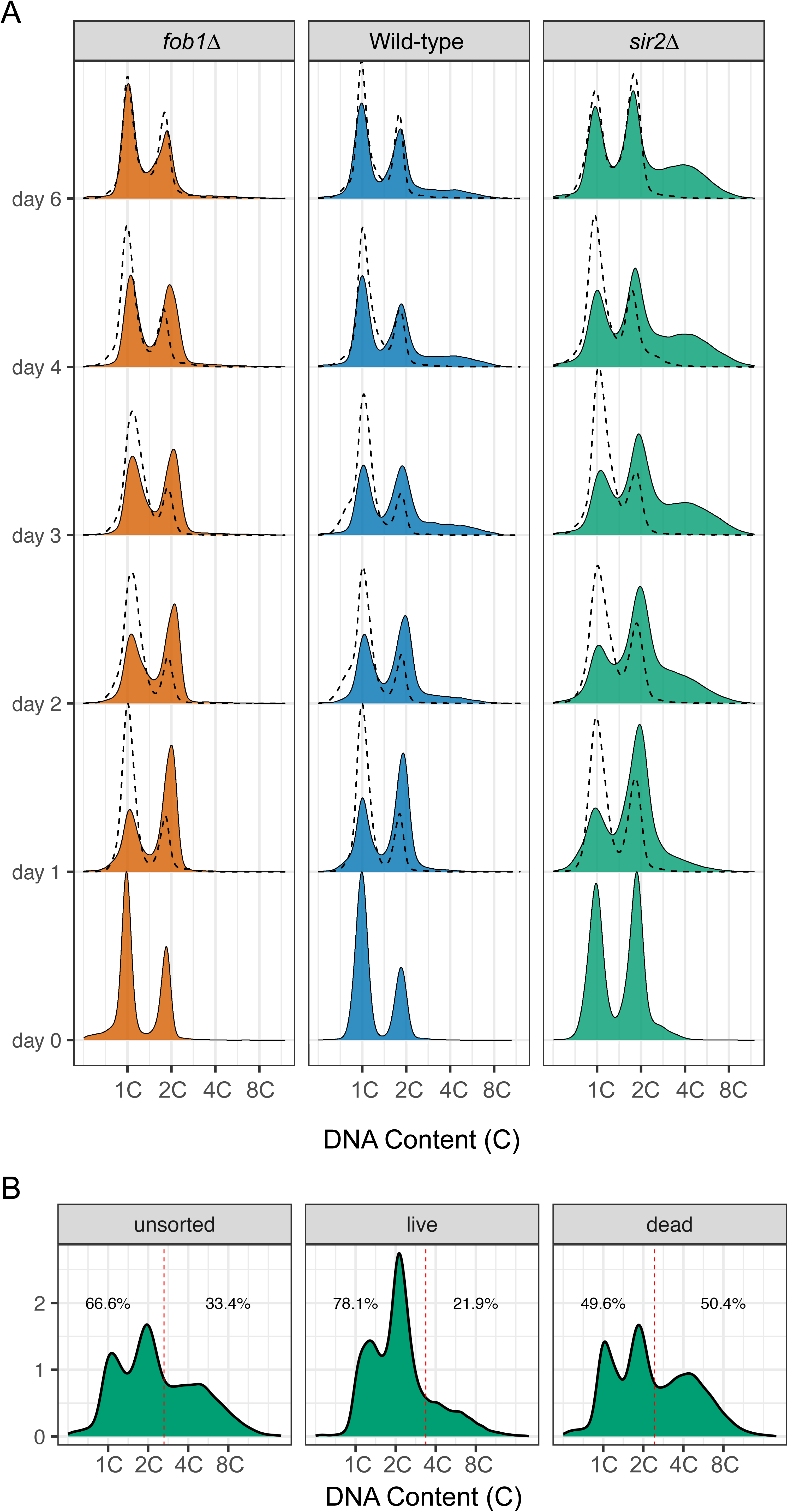
rDNA accumulation is heterogenous in replicate 2 of the aging populations. See the legend to Fig 8 A and D for a description of the flow cytometric analyses of the three population of replicate 2 and the live/dead sorted population of day 4 *sir2Δ* mother cells.

**Supplementary Figure 5:**
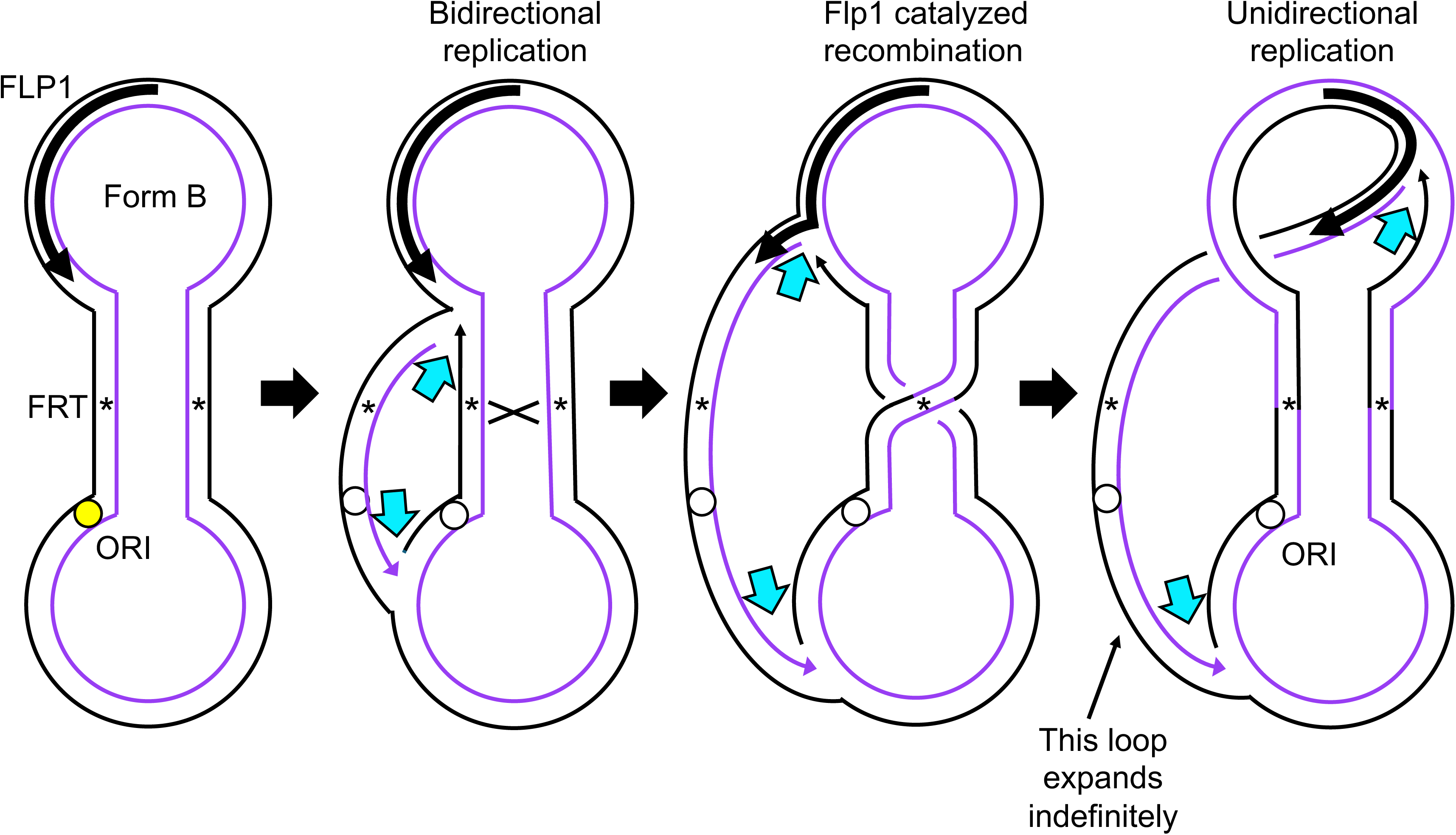
Futcher model for 2-micron amplification. The 6.3 kb 2-micron plasmid contains two 599 bp inverted repeats that partition the plasmid into two unique loops—one that contains the origin of replication (ORI) and the other that contains the *FLP1* gene. The plasmid exists in two isomeric forms (A and B; form B is shown at the left) that differ in the orientation of one loop with respect to the other. Their interconversion is catalyzed by the site-specific recombination activity of Flp1 that recognizes the FRT site in each repeat. When recombination between inverted repeats of differing replication status occurs, the two forks that would normally meet at a site opposite the origin to terminate replication, are now moving in the same direction around the 2-micron plasmid. As they chase each other around the plasmid, they generate a loop of increasing length that is composed of multimers of the 2-micron plasmid. The Futcher model (1986) was the first example where recombination between repeats of different replication status was proposed to lead to DNA amplification. Images are redrawn from Futcher (1986) and show the two strands before and after the recombination event.

**Supplementary Figure 6:**
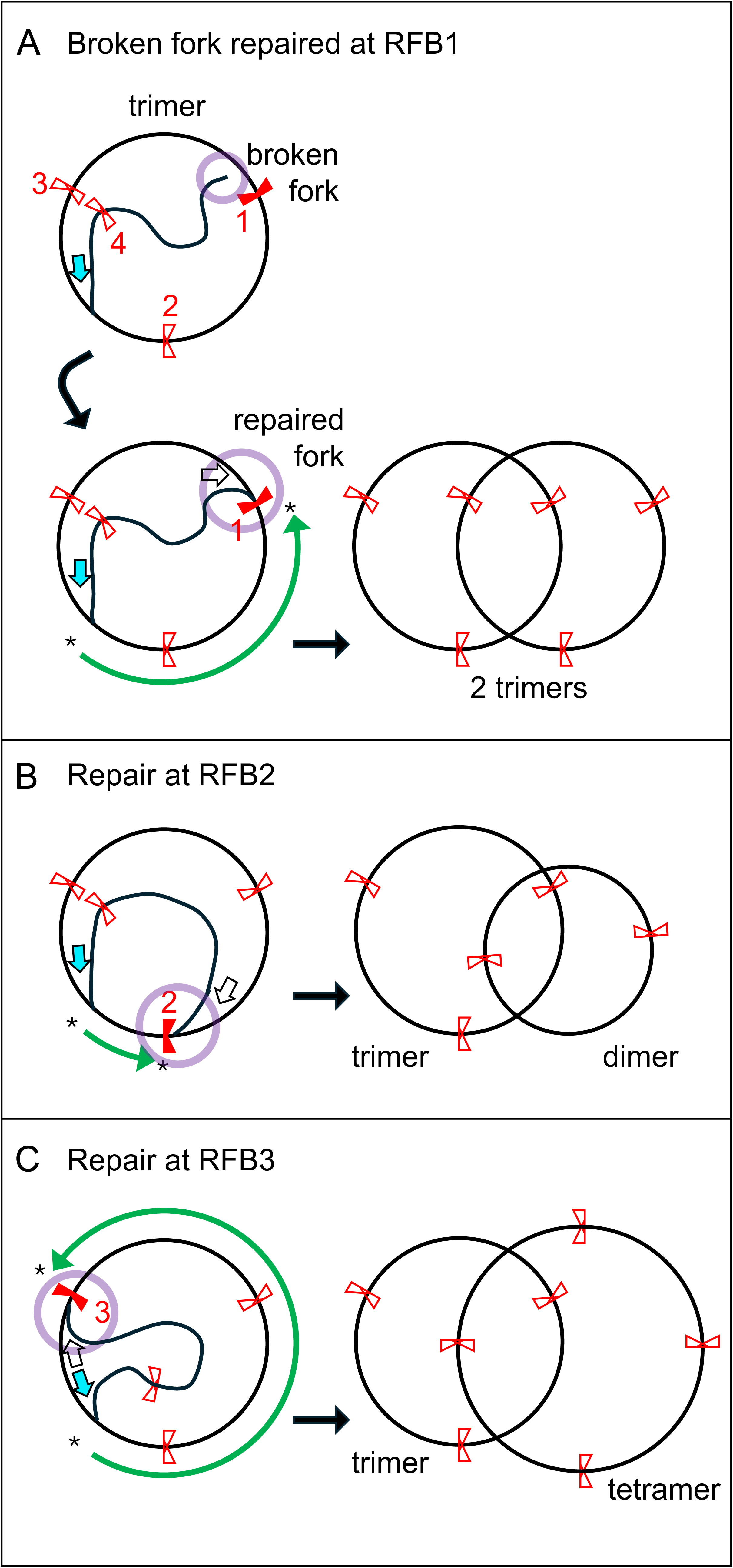
Intramolecular repair of an ERC with a broken fork. A) A trimer ERC in the process of replication suffers a broken fork at RFB1. Repair using the repeat adjacent to RFB1 reestablishes bidirectional forks (cyan and white arrows) and after completion of replication (green curved arrow) produces two trimer ERCs. B and C) The same broken fork adjacent to RFB-1 can repair using the repeats next to RFB-2 or -3 and generates two ERCs after completion of replication. One of the sister molecules remains a trimer; however, the other sister molecule has either two or four repeats, respectively.

**Supplementary Figure 7:**
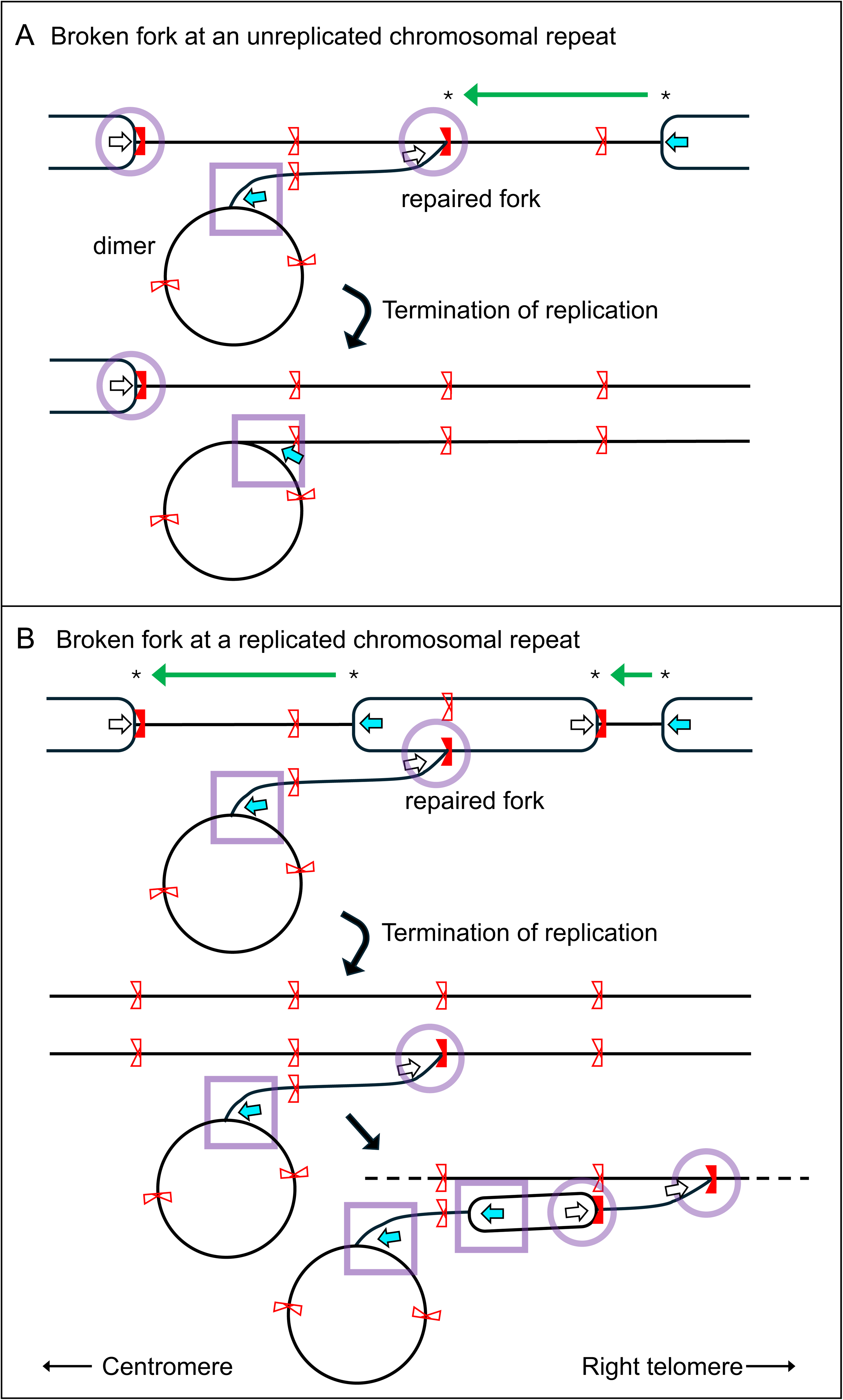
Repair of an ERC with a broken fork using a chromosomal rDNA repeat. Illustrated is a dimer ERC with a broken fork using a chromosomal rDNA repeat for repair. A) If the repeat used for repair is unreplicated, then after all correctly oriented forks have terminated (green arrows) the two sister chromatids will contain branches and be genetically unbalanced—one chromatid has two copies of sequences to the left of the rDNA including the duplicated centromeres and the other is missing a centromere and all sequences to the left of the rDNA. B) If the repair template has already been replicated then one of chromatids will have a side branch of rDNA repeats ending in a lariat with a runaway (rolling circle) fork. In subsequent cell cycles, replication initiation in the branch that connects the ERC to the chromosome adds additional runaway and blocked forks while the branch continues to lengthen from the original runaway fork on the ERC. The branched chromosomes in A and B would remain in the well of a CHEF gel and their runaway forks could expand rDNA hybridization signal in the well.

**Supplementary Figure 8:**
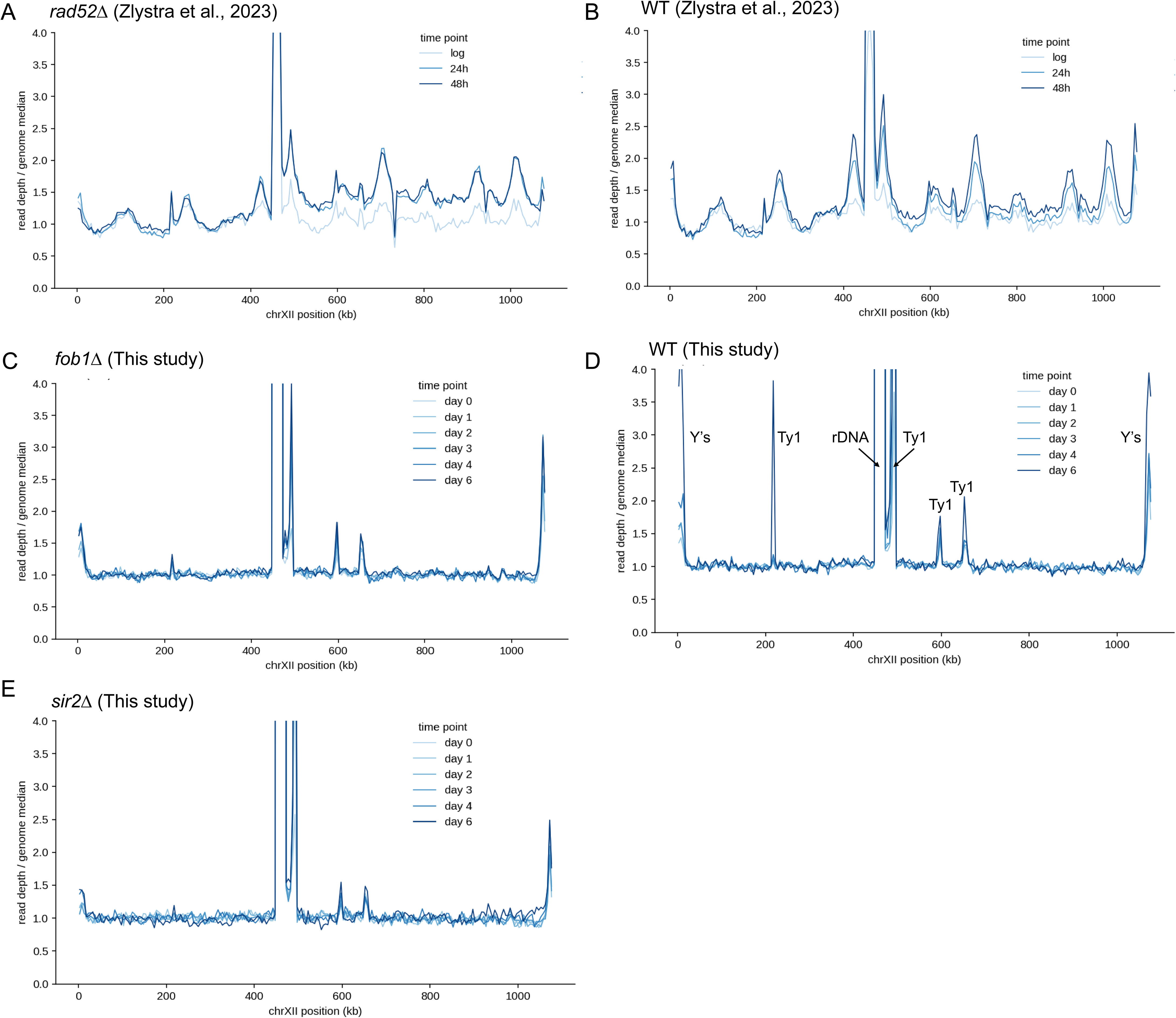
Whole genome sequencing of aging populations. A) Normalized read depth for Chr XII in a *rad52Δ* mutant from Zlystra et al. (2023) in log phase and after 24 and 48 hours of aging. B) Normalized read depth of Chr XII in a wild-type (WT) aging population from Zlystra et al. (2023) across the same sample times as in (A). C-E) Normalized read depth for Chr XII in *fob1Δ*, WT and *sir2Δ* populations, respectively, for replicate 2 from this study. Repeated features (telomeric Y’s, transposable elements Ty1 and the rDNA locus) are labeled in the WT panel.

